# Long-term effects of an elephant-dominated browser community on the architecture of trees in a fenced reserve

**DOI:** 10.1101/2024.12.18.629108

**Authors:** Lucie Thel, Dietre Stols, Sarah Orth, D.D. Georgette Lagendijk, Rob Slotow, Jan A. Venter, Herve Fritz

**Author notes:** Corresponding authors Herve Fritz,; Lucie Thel,; Dietre Stols. These authors share first authorship (Lucie Thel, Dietre Stols). These authors co-supervised the project (Jan A. Venter, Herve Fritz).

## Abstract

African elephants (*Loxodonta africana*), in conjunction with the community of browser species, exert substantial top-down control over the woody vegetation in savannas by utilising large amounts of plant biomass, as well as through non-consumptive effects. However, how much browsers affect the pattern of proportional growth between different tree components remains understudied. Using vegetation data collected in 2000-2001 and 2019 for more than 3,500 trees inside and outside Madikwe Game Reserve, South Africa, we determined the long-term effects of an increasing elephant population, in conjunction with the community of meso-browsers, on structural relationships in 13 tree species. The number of trees utilised by elephants increased between 2000 and 2019, but individuals were not more intensively utilised. After almost two decades of use by elephants, we observed a reduction in the logged initial growth rate of the structural relationship between tree height and stem diameter, without modification of the asymptotic change in growth rate. Despite species-specific variability, tree height was overall reduced for a given stem diameter. Canopy area, as well as its structural relationship with stem diameter remained mostly stable. We suggest that elephants are responsible for hedging by reducing tree height. Together with impala (*Aepyceros melampus*), the dominant species in this meso-browser community, they could stimulate regrowth by browsing the canopy of the vegetation maintained in the browsing trap. Our study emphasises the necessity of long-term, species-specific studies to improve our understanding of how the browser community, and elephant in particular, affect structural relationships in trees.

## 1. INTRODUCTION

Savannas, which cover approximately half of sub-Saharan Africa (du Toit and Cumming, 1999) and a fifth of the earth’s land surface (Sankaran et al., 2005), are a dominant biome characterised by a continuous grass understory interspersed by a discontinuous woody overstory (Scholes and Archer, 1997). The productivity of these savannas is shaped by a complex interplay of rainfall, soil dynamics, micro- and macro-herbivory, as well as fire and frost regimes, which contribute to the diversity of plant species and the structure of the vegetation in these landscapes (Scholes and Archer, 1997; Du Toit and Cumming, 1999; Van Langevelde et al., 2003; Sankaran et al., 2005; Sankaran et al., 2008; Holdo et al., 2009; Shannon et al., 2011). Although fire is commonly considered the dominant factor shaping savanna vegetation by reducing canopy cover and maintaining a continuous grass understory competing with trees, the herbivore community also strongly influences both vegetation structure and plant community composition through the consumption of biomass (Bond et al., 2005; Danell et al., 2006; Sankaran et al., 2008; Lagendijk et al., 2015). Herbivory can theoretically lead to a decrease of up to 30 % in tree abundance and 57 % in grass abundance (Staver et al., 2021).

In African savannas, historically characterised by a remarkably high large herbivore biomass (Hempson et al., 2015), the woody vegetation is influenced by a rich community of browsers (Sankaran et al., 2013; Staver and Bond, 2014). Among them, the African elephant (*Loxodonta africana*) plays a determinant role as an ecosystem engineer (Jones et al., 1994; Cumming et al., 1997; Valeix et al., 2011). Elephants affect species diversity by reducing the density of palatable tree species (e.g., *Senegalia mellifera*, *Sclerocarya birrea caffra*; Sankaran et al., 2013; Cook et al., 2017), sometimes causing local extirpation of highly selected species (e.g., baobab *Adansonia digitata* and *Commiphora ugogensis*; Barnes, 1983; O’Connor et al., 2007). Although elephants are known for pushing over, uprooting and debarking trees, which can cause mortality, they also consume biomass, topple stems and break branches, leaving the trees alive but modifying their architecture (O’Connor et al., 2007; Boundja and Midgley, 2010; Cook et al., 2017).

Tree allometry (i.e., the proportional relationships between different components of an individual through growth; Huxley and Teissier, 1936) is the result of the interaction between abiotic factors such as soil composition, water availability, fire frequency, and browsing pressure (Archibald and Bond, 2003; Moncrieff et al., 2011). To persist despite browsing, tree species have evolved various mechanisms of compensation (e.g., coppicing in *Colophospermum mopane*; Lewis, 1991) or protection (e.g., spinescence or cage effect in *Ziziphus mucronata*; Charles-Dominique et al., 2017). Some of these evolutionary modifications, as well as the phenotypic changes imposed by direct removal of biomass by browsers, and in particular elephants, lead to changes in tree allometric and structural relationships (e.g., ratio between tree height, canopy volume, shoot density and total basal area; Styles and Skinner, 2000; MacGregor and O’Connor, 2004; Makhabu et al., 2006; Mapaure and Moe, 2009; Ferry et al., 2021; O’Connor et al., 2024).

The structural modifications generated by elephants in particular, can have cascading impacts on community interactions, such as the modification of forage quality and availability for other browsers (Styles and Skinner, 2000), habitat structure and selection by other mammal and bird species (Valeix et al., 2011; Tripathi et al., 2019), and even biogeochemical processes such as carbon cycle and storage (Sandhage-Hofmann et al., 2021). While earlier studies suggested that elephant activity could transform woodlands and forests into grasslands or open savannas across Africa, a concern known as “the elephant problem” (Laws, 1970; Caughley, 1976), more recent research indicates that the effect of elephants on savanna vegetation is highly variable. A meta-analysis highlighted that these effects depend on environmental conditions, such as rainfall and geographical constraints (Guldemond and Van Aarde, 2008). Additionally, the expected changes might be less striking in open systems than in fenced reserves sustaining high population densities, or when measured over longer periods (Skarpe et al., 2004; Guldemond and Van Aarde, 2008; Ferry et al., 2021).

In this study, we determined the effect that elephants, in conjunction with the meso-browser community, have on structural relationships in 13 dominant tree species. We used historical data from woody vegetation monitoring conducted in 2000-2001 and repeated in 2019 in Madikwe Game Reserve (hereafter MGR), which was exposed to an increasing elephant population, and in the adjacent Barokologadi Communal Property (hereafter BCP) in 2019, which did not contain elephants. We examined how the browser community affected the relationship between stem diameter, tree height and canopy area. We expected a decrease in the coefficients of the structural relationship between tree height and stem diameter with increasing elephant density. Because of the ability, and regular habit, of elephants to break the main stems of trees, mature trees with large stem diameters should be shorter as elephant utilisation increases in MGR through time and comparatively to BCP, the elephant free area. Additionally, we expected a decrease in the coefficients of the structural relationship between canopy area and stem diameter. In MGR, meso-browsers should reduce canopy area as elephants catalyse a browsing trap by reducing tree height (Smallie and O’Connor, 2000; Staver and Bond, 2014). We expected species-specific patterns due to elephant selectivity and tree species life history traits.

## 2. METHODS

### 1. Study site and species

MGR (24°49′ S - 26°13′ E, South Africa), is a fenced reserve currently covering 750 km² (Supporting information 1). MGR is situated within the central bushveld bioregion, as part of the broader savanna biome. The dominant soils are dolomites, black clay and red clay loam (Nel, 2022, unpublished report). The reserve usually experiences a dry season from mid-April to mid-November (Mucina and Rutherford, 2006), with an average annual rainfall of approximately 475-520 mm (Trinkel et al., 2010).

In 2001, the elephant population was estimated at 389 individuals (i.e., 0.60 elephants/km^2^; reserve size formerly 650 km^2^; North West Parks and Tourism Board, 2024). By 2019, the elephant population increased almost threefold, with an estimated 1,318 individuals (i.e., 1.76 elephants/km^2^; North West Parks and Tourism Board, 2024). Elephants constituted approximately 70 % of the herbivore biomass in MGR at the time of the study (North West Parks and Tourism Board, 2024). Other browser and mixed-feeder herbivores present in the reserve include giraffe (*Giraffa camelopardalis*), eland (*Taurotragus oryx*), greater kudu (*Tragelaphus strepsiceros*), springbok (*Antidorcas marsupialis*), impala (*Aepyceros melampus*), bushbuck (*Tragelaphus scriptus*) and black rhinoceros (*Diceros bicornis*). Although the meso-herbivore population remained mostly stable, the density of impala increased substantially between 2011 and 2019, from 3.84 individuals/km^2^ to 6.30 individuals/km^2^ (Fig. 1).

**Figure 1:**
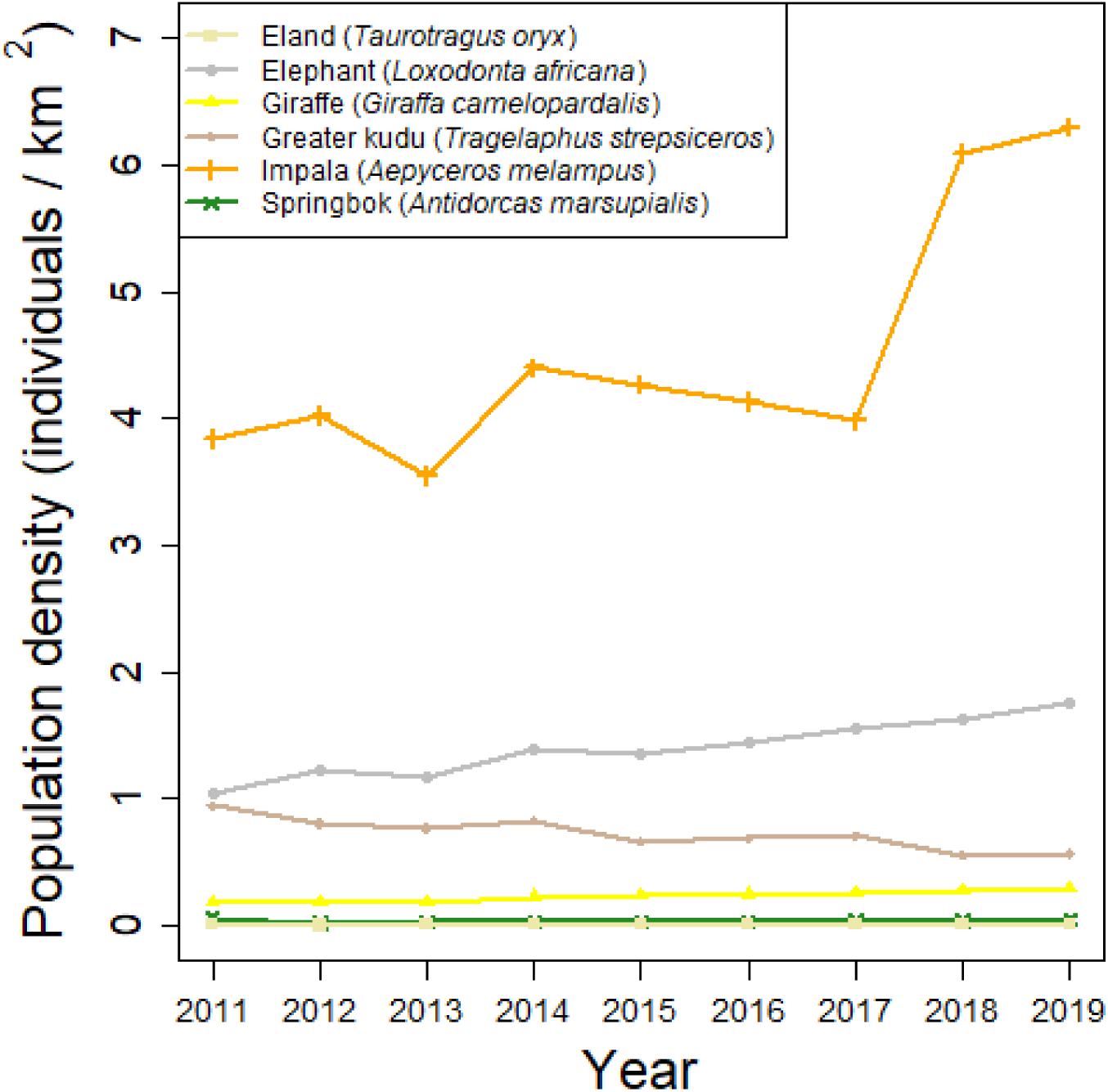
Changes in population density of browser and mixed-feeder species in Madikwe Game Reserve between 2011 and 2019.

BCP covers 160 km^2^ (Supporting information 1) and is separated from MGR by an elephant-proof fence, rendering it an elephant-free zone. The physico-chemical conditions of the area are similar to MGR, and the dominant soils are black clay and red clay loam (Nel, 2022, unpublished report). BCP has been used for cattle, goat, and sheep farming for meat production since 2009, with occasional firewood collection by local communities (Barokologadi Communal Property Association, unpublished report), all of which can affect the structure of the woody vegetation (e.g., Scogings and Macanda, 2005; Tripathi et al., 2019). Bushbuck, greater kudu and impala roam freely in BCP in unknown numbers, but at lower densities than in MGR. Overall, BCP undergoes very little disturbance compared to MGR.

As no fire records were available locally for the study sites, we used the MODIS Burned Area Monthly Global 500 m layer from the NASA website (data retrieved on 2025-03-10 from https://modis.gsfc.nasa.gov; Giglio et al. 2018) to estimate fire occurrence in the study site between 2000 and 2019 (satellite data only available since 2000). Fire should primarily affect tree survival and density (Van Langevelde et al., 2003), but only rarely tree structure. Indeed, young and short individuals would simply be removed by fires, while mature and tall individuals outside the fire trap should remain mainly unaffected (Morrison et al., 2016). Medium individuals are the most susceptible to fire in terms of structure and can be subjected to reduced canopy volume. Nevertheless, such an effect would be noticeable in a relatively short time scale after the last fire occurring in the area. MGR experienced yearly fires between 2001 and 2012 (except in 2007), but fire has been absent since then. BCP seldom experienced fires between 2000 and 2019 (in 2002, 2003, 2006, 2010 and 2011). We thus expected no major effect of fire on observed tree structure in this study (Schafer and Mack, 2014).

### 2. Sampling design

Tree morphological measurements were collected at 35 sites in MGR during three two-weeks periods in January 2000, July 2000 and January 2001, following a stratified random sampling protocol. Sampling sites were selected randomly to cover the different vegetation types present in the study sites (covering mostly three dominant vegetation types: western sandy bushveld, dwaalboom thornveld and Madikwe dolomite bushveld, Mucina et al., 2008). The minimal distance between sites was approximately 100 m. In each site, a 50-meter long transect was used to record woody plants while adjusting width for tree density. The transect was laid out using a central line as the base from which to measure distances to the sides of sample rectangles. Depending on the local vegetation density, the distance from the central line to the sides of the sample rectangles was adjusted from two to five meters, in order to minimize sampling time while maximizing the diversity of tree height and tree species. A targeted sampling was performed in some sites to better accommodate low-abundant dead trees, rare species, or to the contrary overabundant species (e.g. *Dichrostachys cinerea*), extending the sample area to a maximum of 50 m on each side of the central line. However, the inconsistencies in transect sizes only marginally influence our analyses as tree abundance was not the object of the present study.

The 35 original sites from 2000-2001 were re-sampled between February and June 2019 (except for two sites sampled in October), as well as 27 new sites in BCP, following the same methodology. In BCP, 23 sites were sampled during a two-week period in October, and four sites were sampled between April and June (covering mostly three dominant vegetation types: western sandy bushveld, Madikwe dolomite bushveld and Dwarsberg mountain bushveld, Mucina et al., 2008). The minimum distance between sites in BCP was approximately 240 m.

We defined three spatio-temporal conditions: i) “Old”, corresponding to MGR data from 2000-2001 and exposed to a relatively low elephant density (n = 2,318 trees, 0.60 elephants/km^2^), ii) “New”, corresponding to MGR data from 2019 and exposed to a higher elephant density (n = 1,582 trees, 1.76 elephants/km^2^) and iii) “Out”, corresponding to BCP data from 2019 and excluding elephants (n = 1,006 trees, 0.00 elephants/km^2^). Although not completely untouched by meso-browsers (as well as occasional firewood collection), the “Out” condition provides a good control condition in the absence of any elephant pressure in a similar landscape.

For each tree sampled, we recorded the number of live stems and measured stem diameters. Diameters were measured with callipers to the nearest 0.10 cm, at 0.50 - 0.75 m above the ground for tall trees (approx. > 2.00 m tall), at 0.10 - 0.50 m above the ground for short trees (approx. between 0.10 and 2.00 m tall), and at ground level for very short trees (approx. < 0.10 m tall). To prevent lengthy measurements in the field in the case of multistem trees (mean ± sd number of stems per tree = 4 ± 5 stems, range = 1 - 68 stems), all the stems coppicing the same year were identified as a clump and one of these stem diameters was measured as a reference. All the stems of the same clump were then allocated to the same diameter value. Regular checks on randomly selected stems of the same clump were done to insure the consistency and validity of the grouping. For the analyses, we used the stems within the six largest stem diameters for a given individual (i.e., all the stems of a similar diameter up to six classes, hereafter called “stem classes”, mean ± sd diameter = 2.60 ± 4.83 cm, range = 0.10 to 75.00 cm).

For trees < 2.00 m tall (more than 80 % of the individuals), we measured tree height to the nearest 0.01 m as the vertical distance from the ground to the tree’s highest point, and height of the bottom of the canopy (HBC) as the vertical distance from the ground to the height where the main canopy starts, using a measuring tape. For trees > 2.00 m in height, tree height and HBC were estimated to the nearest 0.50 m using an object of known height as a reference (e.g. measuring pole or one of the field technicians) to facilitate estimation of the height of the tree, as described in Shannon et al. (2008).

From these field measurements, we then calculated the total stem diameter as the diameter of a disk of area equal to the sum of all the stem areas, each stem being modelled as a perfect cylinder, as explained in equation (1):

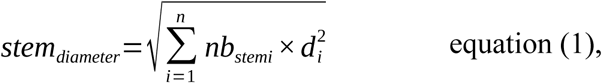

where *d_i_* is the diameter of the stem class *i* and *nb stem_i_* is the number of stems belonging to the stem class *i*. We calculated the vertical area of the canopy, a proxy of the canopy volume taking advantage of the measurements available in the historical data collection protocol, using the formula of the surface of an oval as explained in equation (2):

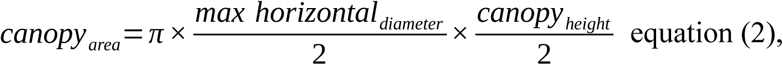

where *max horizontal diameter* corresponds to the widest horizontal diameter of the oval formed by the canopy, and *canopy height* corresponds to the difference between tree height and HBC and represents the vertical diameter of the oval formed by the canopy. A Spearman correlation test confirmed the existence of a high rank correlation between *max horizontal diameter* and *canopy height* (all P < 0.05 except one: *Senegalia mellifera* in the “New” condition: P = 0.08, see Supporting information 2 for details).

To disentangle the influence of meso-(i.e., mostly bushbuck, eland, greater kudu, impala and springbok) and mega-(i.e., mostly elephant and giraffe) browsers in this landscape, we defined three tree height classes following common thresholds used in the current literature (Makhabu, 2005; Sankaran et al., 2013; Staver and Bond, 2014; Lagendijk et al., 2015; Ferry et al., 2021; Thompson et al., 2022): i) short trees, < 0.50 m tall (mainly seedlings and saplings) mostly utilised by meso-browsers, ii) medium trees, between 0.50 m and 2.00 m potentially utilised by both meso- and mega-browsers, iii) tall trees, > 2.00 m (mainly mature trees) utilised by mega-browsers.

For each individual, we estimated an elephant utilisation index as the percentage of biomass (i.e., branches, leaves, stems, bark and/or roots) removed by elephants, using eight categories: 0 (0 %), 1 (1 to 10 %), 2 (11 to 25 %), 3 (26 to 50 %), 4 (51 to 75 %), 5 (76 to 90 %), 6 (91 to 99 %), and 7 (100 %, Walker, 1976). Contrary to meso-browsers, elephant utilisation is easy to discriminate accurately and rapidly in the field due to the characteristic marks left on the trees browsed (Thompson et al., 2022), explaining why only elephant utilisation was recorded in the historical protocol.

### 3. Statistical analyses

All statistical analyses were conducted using R (version 4.4.2, R Core Development Team 2024). We first identified the species that were present in all three spatio-temporal conditions with sufficient sample size (i.e., ≥ 15 trees in each spatio-temporal condition) and included only these species in all our analyses to ensure comparability between spatio-temporal conditions and statistical robustness (n = 13 species; n_Old_ = 1,689 trees, n_New_ = 1,391 trees, n_Out_ = 712 trees; Supporting information 3). In other words, this study focussed on the dominant woody species in this landscape.

We first studied the change in elephant utilisation frequency as the frequency of utilised (i.e. “1” in the model) versus non-utilised (i.e. “0” in the model) individual in “Old” versus “New” conditions using a Generalised Linear Mixed Model (GLMM, Bates et al., 2015) with a binomial distribution of error and a logit link, as well as species and sampling site as random intercepts.

We then investigated the effect of spatio-temporal condition on tree height and canopy area (log-transformed with a natural log to meet assumptions of normality and homoscedasticity) using Linear Mixed Models (LMM), with species and sampling site as random intercepts. We conducted the same analysis for each species individually to check for the presence of species-specific patterns (Supporting information 4). We also tested the correlation between tree height and canopy area using a Spearman’s correlation for each species in each condition, as these two variables are known to be correlated (e.g., Archibald and Bond, 2003; Arzai and Aliyu, 2010; Supporting information 2). We subsequently investigated the effect of tree height class and spatio-temporal condition on stem diameter and canopy area (log-transformed with a natural log to meet assumptions of normality and homoscedasticity) using LMM, with species and sampling site as random intercepts.

To study the structural relationships in trees, we determined the species-specific coefficients of the structural relationship between tree height and stem diameter, and between canopy area and stem diameter (as well as between max horizontal diameter of the canopy and stem diameter, see Supporting information 5), in each spatio-temporal condition. We used stem diameter as a reference in the structural relationships as browsers are expected to have no effect on this parameter (following Moncrieff et al., 2011; Schafer and Mack, 2014). One commonly expected relationship in general tree allometry (Chave et al., 2005) can be written as equation (3):

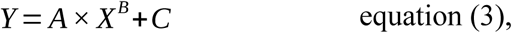

where *X* and *Y* are two tree measurements (*X* is stem diameter and *Y* is either tree height or canopy area here), *A* corresponds to the mathematical term “initial growth rate” of the relationship between *Y* and *X*, *B* corresponds to the “asymptotic change in growth rate” of the relationship, and *C* is a constant close to 0. *C* is 0 in nature as a tree with no stem cannot have a measurable height or canopy. If *C* is significantly different from 0, it suggests poor fit of the model. Mathematically, the initial growth rate *A* illustrates the intrinsic growth of the structural relationship linking the two parameters *X* and *Y*, namely the growth potential that can be reached. The asymptotic change in growth rate *B* corresponds to the speed of the increase in growth, namely how fast the curve increases. If *B* < 1 the growth rate in tree height (canopy area, respectively) decreases as the stem diameter increases, if *B* > 1 the growth rate increases with increasing stem diameter.

However, in our study, visual inspection revealed non-normality and heteroscedasticity associated with an important variance in tree height and canopy area, creating a poor fit of the data with equation (3). Therefore, we used the log-transformed formula (under the assumption of C = 0), and we fitted the corresponding linear regression instead, as described in equation (4):

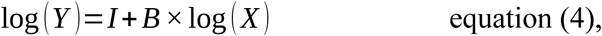

where *X* and *Y* are the same two tree measurements, *B* remains the asymptotic change in growth rate described in equation (3), and *I* corresponds to the logged initial growth rate *A* described in equation (3). In each spatio-temporal condition, we fitted such a relationship for each species separately. We tested the effect of sampling site as a random intercept with the “ranef” function of the “lme4” package (Bates et al., 2015). We removed it when it had no effect and prevented the convergence of the model (n = 6 models out of 39 for both tree height and canopy area relationships, see details in Supporting information 5). We extracted the coefficients *B* and *I* for each species in each spatio-temporal condition, and we assessed the effect of the spatio-temporal condition on the variability of these coefficients using a one-way ANOVA followed by Tukey post-hoc pairwise comparison tests.

We then reconstructed the initial structural relationships between tree height (canopy area, respectively) and stem diameter as described in equation (3), for each spatio-temporal condition. We used the average of the coefficients *B* and *I* estimated for each species with equation (4). We weighted these averages using species frequencies in each spatio-temporal condition to account for the relative importance of each species in the overall relationship. To assess the quality of the fit, we calculated the residuals of this fit, namely the difference between the observed and the predicted values, for each tree height class (Supporting information 6).

## 3. RESULTS

We identified 13 species represented by more than 15 individuals in all three spatio-temporal conditions: *Combretum hereroense*, *Dichrostachys cinerea*, *Euclea undulata*, *Flueggea virosa*, *Grewia flava*, *Grewia monticola*, *Gymnosporia buxifolia*, *Senegalia erubescens*, *Senegalia mellifera*, *Vachellia karroo*, *Vachellia tortilis*, *Ximenia americana* and *Ziziphus mucronata* (Supporting information 3). The most abundant species within all three spatio-temporal conditions was *Dichrostachys cinerea* (n_old_ = 327, n_new_ = 482, n_out_ = 193), and the least abundant species was *Senegalia mellifera* in the “Old” and “New” conditions (n_old_ = 23, n_new_ = 15, n_out_ = 67), as well as *Vachellia karroo* in the “Out” condition (n_old_ = 108, n_new_ = 54, n_out_ = 16). Both MGR and BCP were characterised by shrub-dominated vegetation, with more than 80 % of the trees < 2.00 m tall (1.24 ± 1.29 m tall on average, Fig. 2a) and an average canopy area of approximatively 1.80 ± 4.46 m^2^ (Fig. 2b).

**Figure 2:**
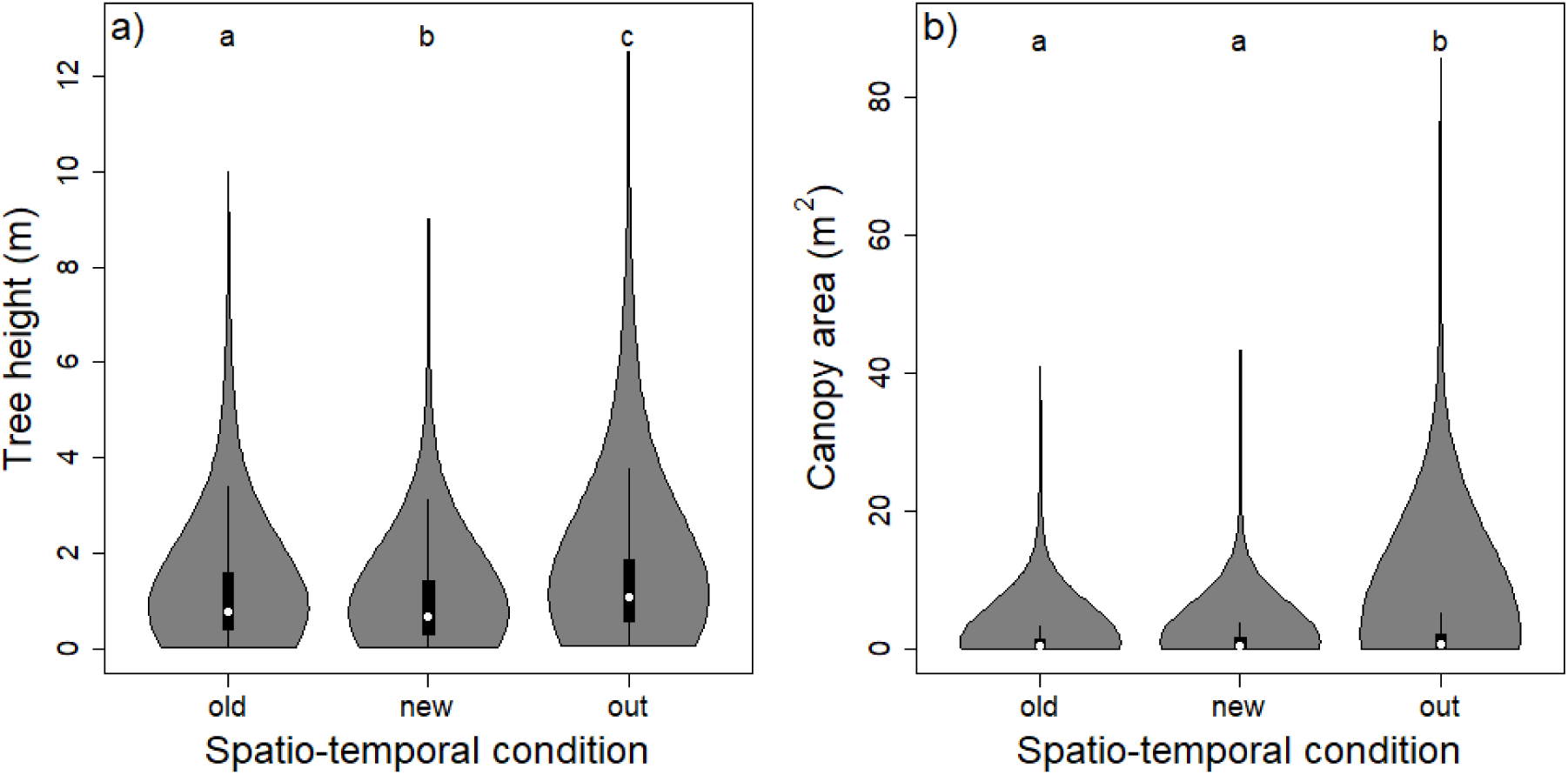
Variation of a) tree height and b) canopy area, between the three spatio-temporal conditions in Madikwe Game Reserve (MGR) and Barokologadi Communal Property (BCP), South Africa. “Old”, data collected in MGR in 2000 and 2001 at relatively low elephant density; “New”, data collected in MGR in 2019 at relatively high elephant density; “Out”, data collected in BCP in 2019 in the absence of elephants. All 13 tree species were pooled. White dots correspond to the median, black boxes correspond to the interquartile range (IQR) of the distribution, whiskers extend to the minimum and maximum values within 1.5 times the IQR and shaded areas correspond to the kernel density estimation of the distribution.

Elephant utilisation varied between 0 and 7 in all tree height classes (Supporting information 7), both in “Old” and “New” conditions (except for the fact that none of the shortest and tallest trees in “Old” were completely utilised, Table 1). Most of the trees in our dataset were not utilised at all by elephants (90 and 60 % of non-utilised trees in the “Old” and “New” conditions, respectively), but there was a significant increase in the amount of trees utilised between “Old” and “New” (beta = 1.91 ± 0.11, P < 0.01, P_utilised-Old_ = 0.07 [0.01; 0.35], P_utilised-New_ = 0.33 [0.07; 0.78]). *Gymnosporia buxifolia*, *Senegalia erubescens* and *Vachellia tortilis* were the most utilised species in “Old” (between 20 and 30 % of the trees utilised), while *Ximenia americana* was not utilised at all. More than 20 % of the trees of all 13 species, except from *Ximenia americana* and *Euclea undulata*, were utilised at various degrees in “New”. At least 50 % of the trees of *Senegalia erubescens* and *Grewia flava* were utilised to various degrees.

**Table 1:** Number of trees per level of elephant utilisation index for each tree height class in Madikwe Game Reserve (MGR) spatio-temporal conditions: “Old”, data collected in MGR in 2000 and 2001 at relatively low elephant use (n_old_ trees = 1689); “New”, data collected in MGR in 2019 at relatively high elephant use (n_new_ trees = 1391).

| Tree height class | < 0.5 m |  | 0.5 to 2 m |  | > 2 m |  |
| --- | --- | --- | --- | --- | --- | --- |
| Elephant utilisation index | old | new | old | new | old | new |
| 0 | 496 | 426 | 808 | 356 | 199 | 56 |
| 1 | 3 | 21 | 5 | 61 | 9 | 34 |
| 2 | 3 | 17 | 32 | 55 | 34 | 29 |
| 3 | 4 | 18 | 17 | 80 | 38 | 41 |
| 4 | 8 | 23 | 6 | 67 | 11 | 11 |
| 5 | 3 | 35 | 6 | 23 | 2 | 7 |
| 6 | 1 | 10 | 2 | 9 | 1 | 1 |
| 7 | 0 | 1 | 1 | 6 | 0 | 4 |

Overall, trees were significantly shorter in “New” than in “Old” (P < 0.01) and in “New” than in “Out” (P < 0.01, Fig. 2a) conditions. The canopy area was similar in “New” and “Old” conditions (P = 0.05), but it was significantly larger in “Out” than in “New” conditions (P < 0.01, Fig. 2b). Although these results were mostly consistent among seven species (*Combretum hereroense*, *Dichrostachys cinerea*, *Grewia flava*, *Grewia monticola*, *Vachellia karroo*, *Vachellia tortillis* and *Ziziphus mucronata*), there were species-specific patterns (see Supporting information 4 for details). Notably, *Flueggea virosa* increased in tree height and canopy area between “Old” and “New” conditions, and the differences in tree height and canopy area were less pronounced between “New” and “Out” for *Gymnosporia buxifolia*. No significant differences in canopy area were detected between spatio-temporal conditions for *Senegalia erubescens*, *Senegalia mellifera*, *Ximenia americana*. We found a significant and very high rank correlation between tree height and canopy area in all species in all three spatio-temporal conditions (Spearman test: all ρ > 0.6, all P < 0.01, Supporting information 2). Overall, stem diameters were significantly different from each other across spatio-temporal conditions and tree height classes, except in some instances. Notably, we did not detect significant differences between the tallest trees in “Out” and “New” conditions, and the shortest and medium trees in “New” and “Out” conditions (Fig. 3a, see details in Supporting information 8). Canopy areas were also mostly, but not always significantly different from each other across spatio-temporal conditions and tree height classes. Notably, we found no significant difference between the shortest trees in all three conditions, between the medium trees in “Out” and “New” conditions, and between the tallest trees in “Old” and “New” conditions (Fig. 3b, see details in Supporting information 8).

**Figure 3:**
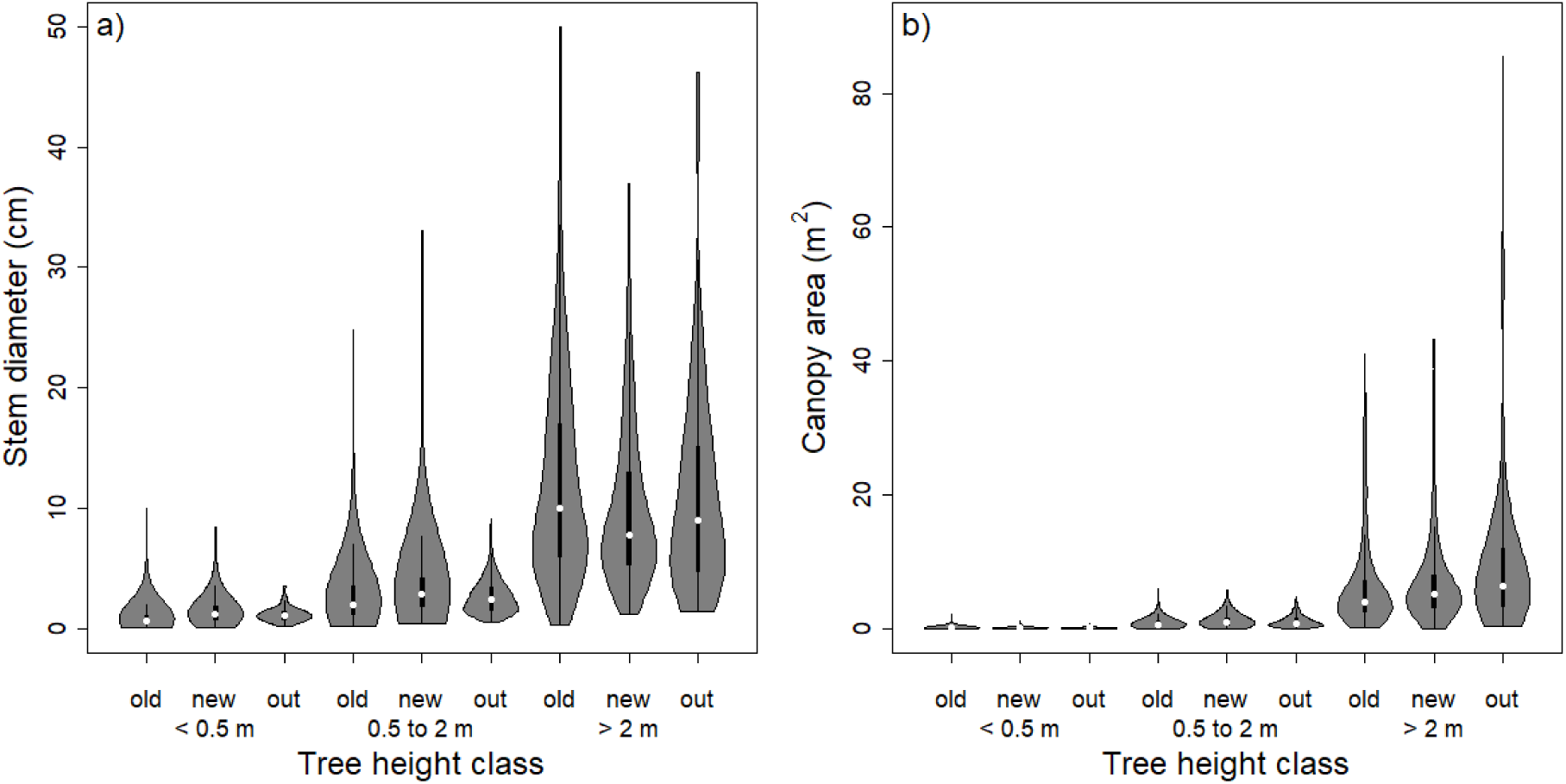
Distribution of: a) stem diameters and b) canopy areas, according to tree height classes (< 0.5 m, from 0.5 to 2 m, > 2 m) in the three spatio-temporal conditions: “Old”, data collected in Madike Game Reserve (MGR) in 2000 and 2001 at relatively low elephant density; “New”, data collected in MGR in 2019 at relatively high elephant density; “Out”, data collected in Barokologadi Communal Property (BCP) in 2019 in the absence of elephants. White dots correspond to the median, black boxes correspond to the interquartile range (IQR) of the distribution, whiskers extend to the minimum and maximum values within 1.5 times the IQR and shaded areas correspond to the kernel density estimation of the distribution.

Regarding the structural relationships, we found similar positive trends between stem diameter and tree height (canopy area, respectively) for all 13 species in all three spatio-temporal conditions (note species-specific variability related to the spatio-temporal condition, such as in *Grewia monticola*, and a negative relationship for *Senegalia mellifera* in the “New” condition, Supporting information 5). Regarding the coefficients of the structural relationships, *I* was significantly lower in “New” than in “Old” conditions in the tree height - stem diameter relationship (Tukey test: P = 0.02, Fig. 4b). Although not significant (ANOVA: P = 0.12), we found a similar trend for *I* in the canopy area - stem diameter relationship (Fig. 4d). *B* was significantly different according to the condition in the tree height - stem diameter relationship (ANOVA: P = 0.04, Fig. 4a), although no pairwise difference was significant (Tukey: all P > 0.05). *B* was not significantly different between “New” and “Old” (Tukey: P = 0.88), but was close to be between “New” and “Out” (Tukey: P = 0.06) conditions for the canopy area - stem diameter relationship (Fig. 4c). The reconstructions of the structural relationships (equation (3)) based on the averaged coefficients extracted from equation (4) showed a good fit in “Out” condition for both tree height and canopy area (Fig. 5a and 5b, Supporting information 6). The predictions slightly under-estimated tree height and to a lower extent, canopy area for tall trees in “Old” condition (Fig. 5a and 5b, Supporting information 6). A similar but stronger pattern was visible in “New” condition, also affecting medium trees (Fig. 5a and 5b, Supporting information 6).

**Figure 4:**
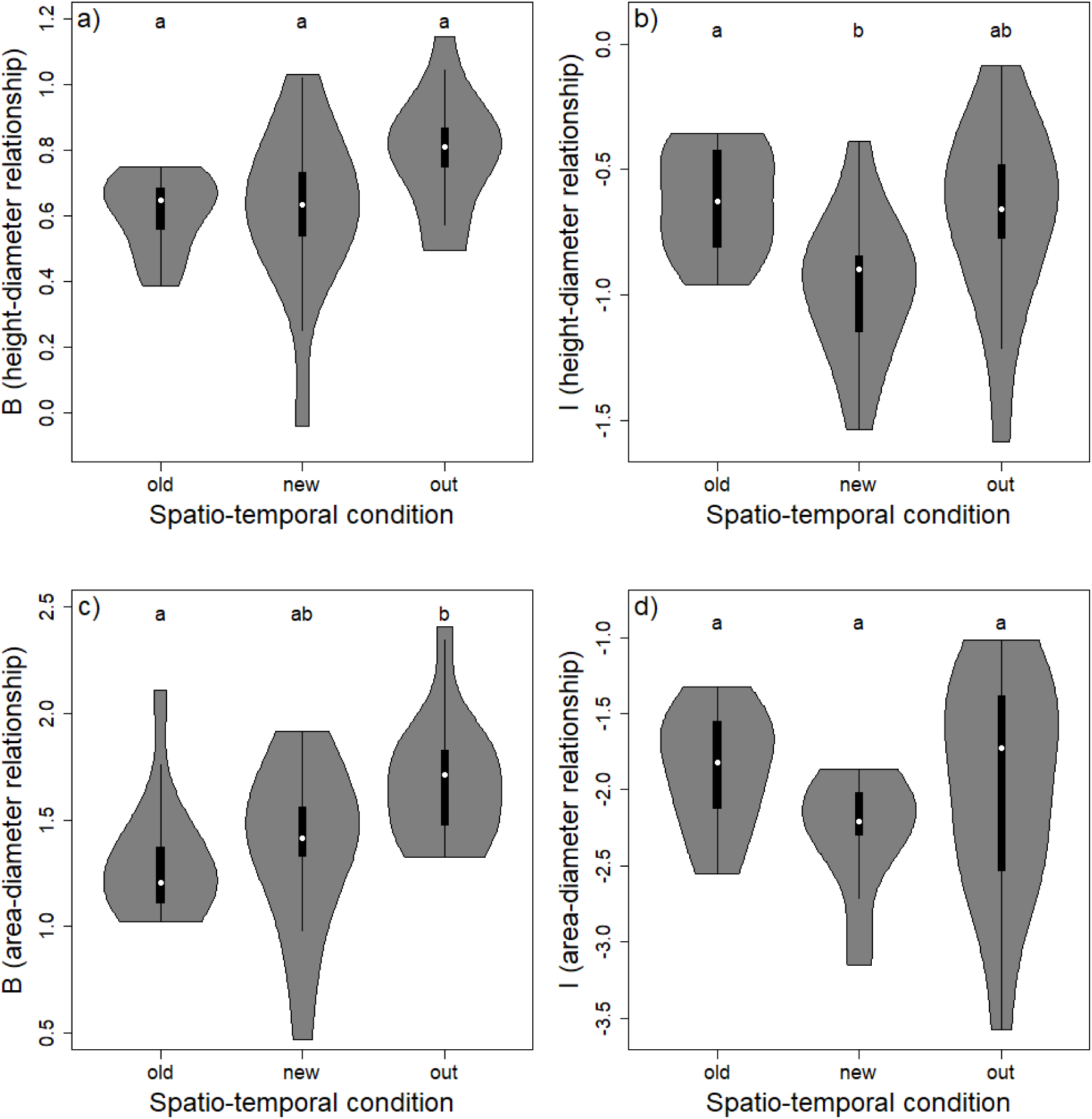
Asymptotic change in growth rate (*B*) and logged initial growth rate (*I*) according to the spatio-temporal condition in Madikwe Game Reserve (MGR) and Barokologadi Communal Property (BCP), South Africa for the tree height - stem diameter and the canopy area - stem diameter relationships: a) *B* for the tree height - stem diameter relationship; b) *I* for the tree height - stem diameter relationship; c) *B* for the canopy area - stem diameter relationship; d) *I* for the canopy area - stem diameter relationship. “Old”: data collected in MGR in 2000 and 2001 at relatively low elephant density; “New”, data collected in MGR in 2019 at relatively high elephant density; “Out”, data collected in BCP in 2019 in the absence of elephants. Letters denote significant differences between spatio-temporal conditions. White dots correspond to the median, black boxes correspond to the interquartile range (IQR) of the distribution, whiskers extend to the minimum and maximum values within 1.5 times the IQR and shaded areas correspond to the kernel density estimation of the distribution.

**Figure 5:**
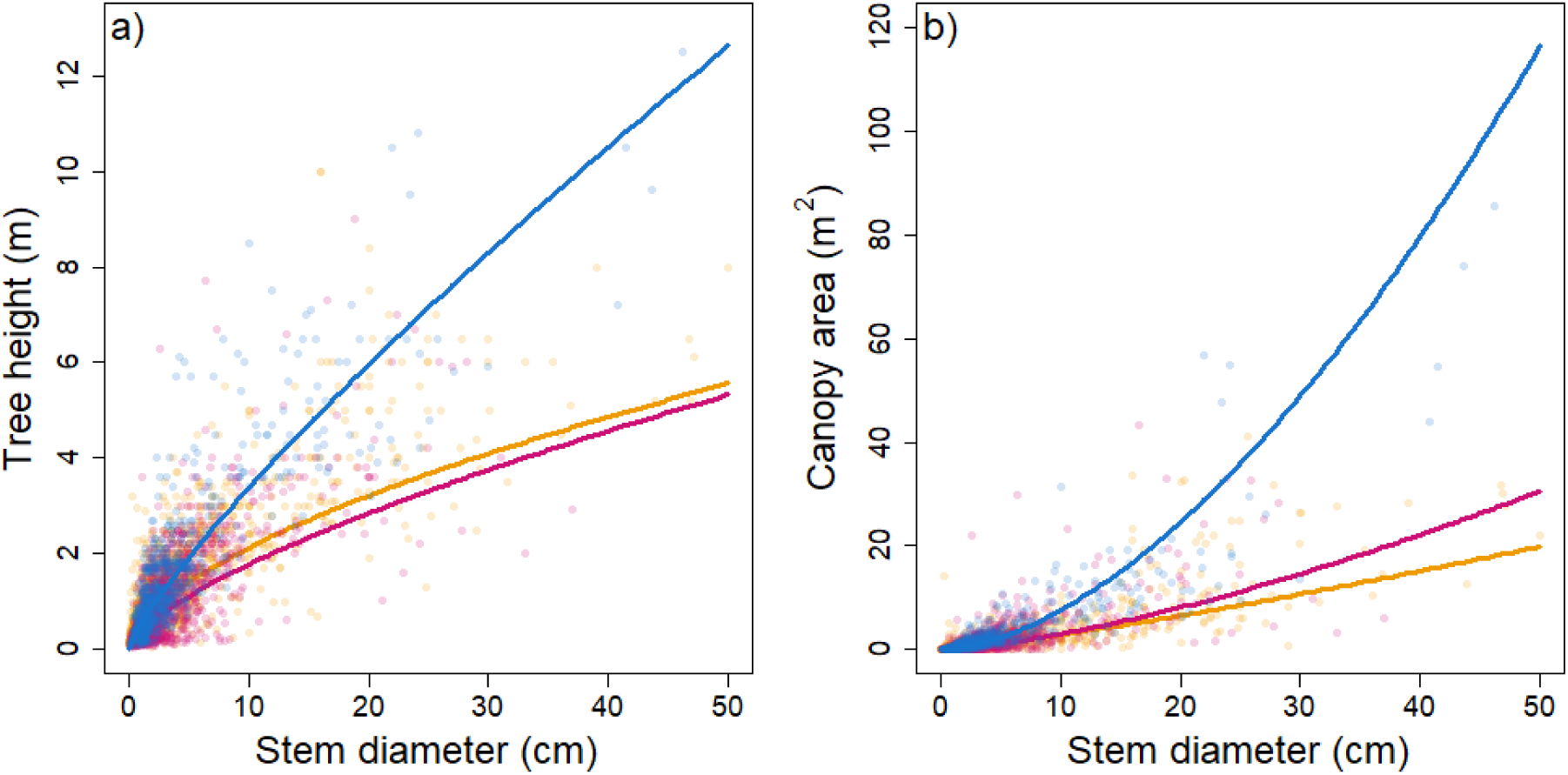
Structural relationship a) between tree height and stem diameter, b) between canopy area and stem diameter, in the three spatio-temporal conditions (“Old”, data collected in Madikwe Game Reserve in 2000 and 2001 at relatively low elephant density; “New”, data collected in MGR in 2019 at relatively high elephant density; “Out”, data collected in Barokologadi Communal Property in 2019 in the absence of elephants). Dots represent the observed data, lines represent the predictions of the model (see text for details).

## 4. DISCUSSION

In this study, we highlighted the effects of an increasing elephant population on the structure of the woody vegetation at the tree community scale. Overall, the number of trees utilised by elephants increased in MGR over time, but most of the trees sampled were not utilised by elephants and the intensity of utilisation remained stable across height classes. More than 80 % of the trees were below 2.00 m tall and thus within the browsing trap (Smallie and O’Connor, 2000; Staver and Bond, 2014). We suggest that the effect of elephants is exacerbated by persistent browsing by meso-browsers. By reducing tree height (known as “hedging”, Styles and Skinner, 2000), elephants create conditions for the woody vegetation to become even more subjected to intense browsing by both elephants and meso-herbivores in a browsing trap, which in turn affect the canopy. Their combined effect generates heterogeneity in tree height and canopy area for a given stem diameter, modifying the structural relationships in trees.

Trees were shorter in MGR in 2019 compared to 2000 (the low elephant density condition) and BCP in 2019 (the elephant-free area). There was a reduction in the logged initial growth rate (*I*) of the structural relationship between tree height and stem diameter, without modification of the asymptotic change in growth rate (*B*) in MGR in 2019 compared to the other conditions, further confirmed by the deceleration of the tree height-stem diameter structural relationship at large diameters in MGR versus BCP. These results denote a reduction of the overall growth potential of trees in MGR, but not of the speed at which the growth potential is reached, and suggests that elephants might participate in the maintenance of short trees in the landscape (e.g., Kalwij et al., 2010) by keeping older tree shorter than expected for their stem diameter. Even at moderate density, elephants are known to reduce the abundance of very large trees in African savannas (e.g., Mapaure and Moe, 2009; Gandiwa et al., 2011; Cook et al., 2017; Watson et al., 2019). Although giraffes can also participate in reducing tree height by consuming the top buds (Makhabu, 2005; Cameron and du Toit, 2007), they are present in very limited numbers in MGR and probably contribute only marginally to the observed effects.

During the study period, impala, which was the dominant species among the meso-browsers, more than doubled in MGR (from 2,879 individuals in 2011 to 6,110 individuals in 2021). Impala, a mixed feeder (Hofmann, 1989), is known to inhibit the recruitment of trees in both the seedling and sapling classes, but also to reduce recruitment into taller height classes as they restrain tree growth within the browsing trap (O’Kane et al., 2012; Sankaran et al., 2013). However, we did not find a significant difference in terms of canopy area in MGR between 2000 and 2019, the canopy area remaining stable or even increasing in short and medium height classes. Although the projection of the structural relationship illustrated a decrease in the canopy area for large diameter trees, the coefficients of the structural relationship remained mostly stable across the three conditions, and there was even a tendency towards an increase in the asymptotic change in growth rate (*B*) in MGR in 2019 versus 2000. These findings, consistent with another study finding no reduction in shrub canopy volume with increasing elephant density (O’Connor et al., 2024), suggest potential compensatory mechanisms in response to browsing.

When exposed to herbivory, trees can develop compensatory mechanisms to balance the loss of biomass (McNaughton, 1983; Paige, 1992; Johnson and Ebersole, 2017), although species-specific patterns are common (*Vachellia tortilis* and *Ziziphus mucronata* showed a significant decrease in canopy area between 2000 and 2019; see also Focardi and Tinelli, 2005). More shoots appear following heavy meso-herbivore browsing (Fornara and du Toit, 2007), and trees broken and toppled by elephant coppice (Lewis, 1991). Alternatively, elephant presence is usually associated with a decrease in tree density (Watson et al., 2019), which might also have reduced competition for light in a dense bushy landscape such as MGR, leading to increased growth of individual canopy, in a similar way that herbivores increase species diversity by reducing light competition in grassland systems (Borer et al., 2014).

The predictions of the structural relationship between tree height (and to a lesser extent, canopy area) and stem diameter were slightly under-estimated in MGR in 2000, and even more so in 2019. To the contrary, they were very accurate in BCP, the relatively undisturbed condition, providing empirical support for the suitability of our approach. The presence of browsers in the system disturbs the expected structural relationship by creating variance in tree height and canopy area for a given stem diameter, and so among every tree height class. The changes observed in the structural relationships in MGR thus originate from a cumulative effect of the heterogeneity generated in small stem diameter trees by all browsers, as well as in large stem diameters utilised only by mega-browsers. Under-estimation in MGR also illustrates the possibility for more complex effects of the browser community on these structural relationships. Moncrieff et al. (2011) found threshold models best described the effect of elephants and giraffes on the same relationship. Here, we adjusted a single regression to describe the structural relationship across all tree height classes, but each browser species acts at its own height (Makhabu, 2005). The effect of a given species might reverberate on different portions of the relationship, even where its effect is not predominant. For instance, impalas affect the shorter trees, such as seedlings and saplings, and thus the mathematical initial growth rate in the relationship, whose fluctuations reverberate on the adjustment at larger stem diameters. Additionally, elephants demonstrate a wide browsing range in terms of tree height (Supporting information 7; Smallie and O’Connor, 2000; Makhabu, 2005), implying that the medium- and short-sized trees are probably affected both by them and the meso-herbivore community.

Each tree species has its own allometry, and reacts differently to browsing, creating a high dispersion in the overall relationship. Some species characterised by higher abundance or palatability can be preferentially selected by herbivores (elephants preferentially select e.g. *Combretum elaeagnoides*, Makhabu, 2005, *Colophospermum mopane*, Smallie and O’Connor 2000, while giraffes select *Acacia* species, Mahenya et al., 2016). In MGR in 2019, *Senegalia erubescens* was the most frequently targeted species by elephants with more than 65 % of the trees utilised, whereas *Euclea undulata* remained mostly untouched (only 6 % of the trees utilised). Depending on the tree and herbivore species, different parts of the tree can be preferentially selected and exposed to various types of modifications. Elephants can topple, strip and break stems, such as in *Vachellia tortilis* (Boundja and Midgley, 2010), utilised in almost 40 % of the cases and with a significantly reduced tree height in MGR in 2019, while giraffes browse exclusively on soft nutritious material such as leaves and shoots (e.g. in *Acacia* species; Mahenya et al., 2016). Such preferences generate the potential for species-specific responses in structural relationships in trees (MacGregor and O’Connor, 2004).

Structural, and more generally, allometric relationships in trees have important evolutionary consequences for their maintenance and reproduction (Styles and Skinner, 2000; Archibald and Bond, 2003). These relationships arise from a complex interplay between tree species traits (e.g. regeneration by regrowth in *Senegalia nigrescens*, Fornara and du Toit, 2007), abiotic factors (e.g. fire regimes; Schafer and Mack, 2014) and herbivore preferences (e.g. different herbivores browse different plant parts depending on their size and diet; Cameron and du Toit, 2007). Understanding the profound effects of these interacting factors on tree structure should be taken to the next step by using technologies such as LiDAR (light detection and ranging) instruments, which can provide a more integrative overview of the three-dimensional structure of trees (e.g., Davies et al., 2018; McNeil et al., 2023).

## Data availability statement

The data and scripts that support the findings of this study are openly available in Zenodo Repository at https://doi.org/10.5281/zenodo.14268021.

## ACKNOWLEDGMENTS

We acknowledge and thank Bruce Page, who designed the sampling protocol, and supervised the sampling in 2000-2001. We thank all University of Kwazulu-Natal students and volunteers involved in the 2000-2001 sampling. We thank the fieldworkers who collected the data in 2019 and managed the data for this study: Yentl Swartz, Chris Brooke, Adriaan Venter, Michelle Marais, Jandre Vermaak, Terry-Lee Honiball. We also thank the North-West Parks and Tourism Board for their logistical support for the study. The 2000-2001 sampling was funded by the Amarula Elephant Research Programme through a donation by Distell (PTY) Ltd, and the University of Kwazulu-Natal, with in-kind support from North West Parks and Tourism Board. Postgraduate Research Scholarship funding from Nelson Mandela University and REHABS funding to DS.

## AUTHORS CONTRIBUTION STATEMENT

**Conceptualization**: LT, DS, JV, HF; **Methodology**: LT, DS, HF; **Formal analysis**: LT; **Investigation**: LT, DS; **Data Curation**: LT, DS, SO, GL, RS, JV; **Writing: Original Draft**: LT, DS, SO; **Writing: Review & Editing**: LT, DS, SO, GL, RS, JV, HF; **Visualisation**: LT; **Resources**: DS; **Funding acquisition**: RS, JV, HF; **Validation**: LT, DS, SO, GL, RS, JV, HF; **Supervision**: JV, HF; **Project administration**: JV, HF.

