## supporting information for "Long-term effects of an elephant-dominated browser community on the architecture of trees in a fenced reserve"

#### Authors details

[1] Department of Conservation Management, Faculty of Science, Nelson Mandela University, George Campus, George, South Africa.

[2] Master de Biologie, École Normale Supérieure de Lyon, Université Claude Bernard Lyon 1, Université de Lyon, 69342 Lyon Cedex 07, France.

[3] Sustainability Research Unit, George Campus, Nelson Mandela University, George, South Africa.

[4] REHABS International Research Laboratory, CNRS-Université Lyon 1-Nelson Mandela University, George Campus, Nelson Mandela University, George, South Africa.

[5] School of Life Sciences, University of KwaZulu-Natal, Private Bag X01, Scottville, 3209 South Africa.

[6] Conservation Ecology Group, Groningen Institute for Evolutionary Life Sciences (GELIFES), University of Groningen, PO Box 11103, 9700 CC Groningen, The Netherlands.

[7] BirdEyes, Centre for Global Ecological Change at the Faculty of Science & Engineering and Campus Fryslân, University of Groningen, Wirdumerdijk 34, 8911 CE Leeuwarden, The Netherlands.

[8] Oppenheimer Fellow in Functional Biodiversity, Centre for Functional Biodiversity, School of Life Science, University of Kwazulu-Natal, Pietermaritzburg 3209, South Africa.

*These authors share first authorship; [#]These authors co-supervised the project

#### Corresponding authors

Herve Fritz,; Lucie Thel,; Dietre Stols,

#### Submission and Acceptance Dates

Received: __________; Revised: __________; Accepted: __________.

#### SUPPORTING INFORMATION

**Supporting information 1**: Location of the sites of data collection in Madikwe Game Reserve (MGR, n = 35) and Barokologadi Communal Property (BCP, n = 27), South Africa (see insert). Grey dots represent the sites in "Old" and "New" spatio-temporal conditions in MGR, blue dots represent the sites in "Out" spatio-temporal condition in BCP.

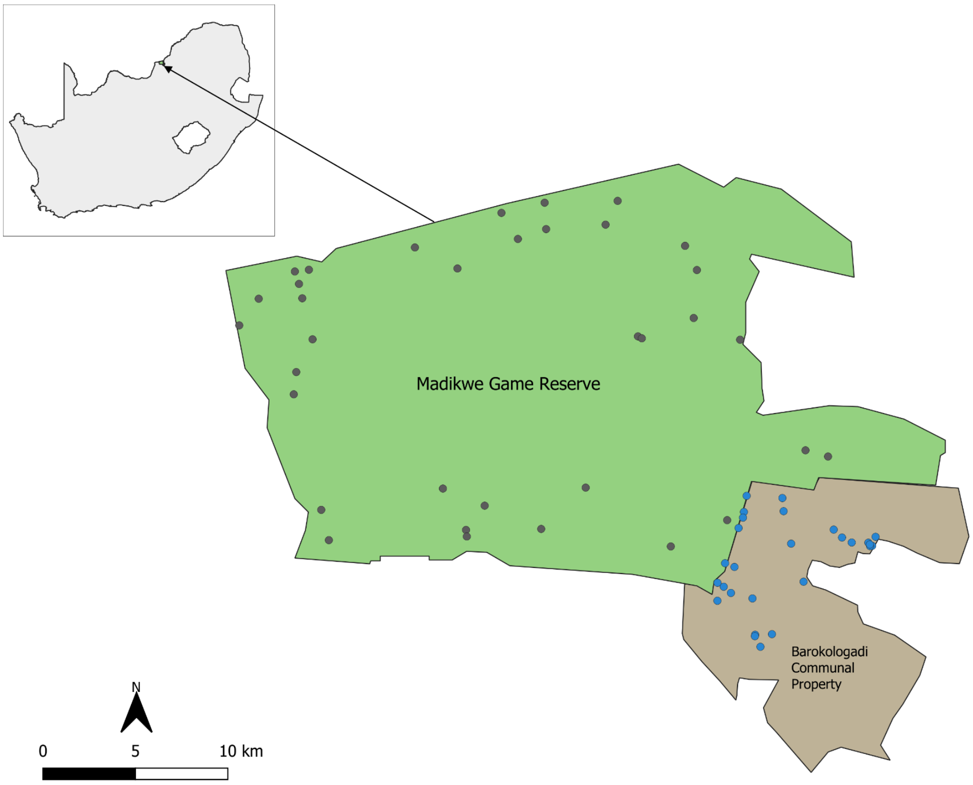

**Supporting information 2**: Species-specific correlations between tree structure parameters.

**Max horizontal diameter and canopy height**

**Table S2.1**: Species-specific Spearman rank correlation (ρ) between max horizontal diameter of the canopy and canopy height, and their significance (P) in Madikwe Game Reserve (MGR) and Barokologadi Communal Property (BCP), South Africa. “Old”: data collected in MGR in 2000 and 2001 at relatively low elephant density; “New”, data collected in MGR in 2019 at relatively high elephant density; “Out”, data collected in BCP in 2019 in the absence of elephants.

| **Species** | $\rho_{old}$ | $P_{old}$ | $\rho_{new}$ | $P_{new}$ | $\rho_{out}$ | $P_{out}$ |
|---|---|---|---|---|---|---|
| *Combretum hereroense* | 0.82 | < 0.01* | 0.84 | < 0.01* | 0.85 | < 0.01* |
| *Dichrostachys cinerea* | 0.76 | < 0.01* | 0.88 | < 0.01* | 0.76 | < 0.01* |
| *Euclea undulata* | 0.70 | < 0.01* | 0.82 | < 0.01* | 0.88 | < 0.01* |
| *Flueggea virosa* | 0.70 | < 0.01* | 0.64 | < 0.01* | 0.63 | 0.01* |
| *Grewia flava* | 0.66 | < 0.01* | 0.79 | < 0.01* | 0.58 | < 0.01* |
| *Grewia monticola* | 0.74 | < 0.01* | 0.83 | < 0.01* | 0.70 | < 0.01* |
| *Gymnosporia buxifolia* | 0.73 | < 0.01* | 0.62 | < 0.01* | 0.43 | 0.03* |
| *Senegalia erubescens* | 0.86 | < 0.01* | 0.87 | < 0.01* | 0.92 | < 0.01* |
| *Senegalia mellifera* | 0.94 | < 0.01* | 0.46 | 0.08 | 0.85 | < 0.01* |
| *Vachellia karroo* | 0.64 | < 0.01* | 0.76 | < 0.01* | 0.75 | < 0.01* |
| *Vachellia tortilis* | 0.86 | < 0.01* | 0.85 | < 0.01* | 0.86 | < 0.01* |
| *Ximenia americana* | 0.44 | < 0.01* | 0.86 | < 0.01* | 0.67 | < 0.01* |
| *Ziziphus mucronata* | 0.83 | < 0.01* | 0.93 | < 0.01* | 0.88 | < 0.01* |

* denotes significance to the level alpha = 0.05.

**Tree height and canopy area**

**Table S2.2**: Species-specific Spearman rank correlation (ρ) between tree height and canopy area and their significance (P) in Madikwe Game Reserve (MGR) and Barokologadi Communal Property (BCP), South Africa. "Old": data collected in MGR in 2000 and 2001 at relatively low elephant density; "New", data collected in MGR in 2019 at relatively high elephant density; "Out", data collected in BCP in 2019 in the absence of elephants.

| **Species** | $\rho_{old}$ | $P_{old}$ | $\rho_{new}$ | $P_{new}$ | $\rho_{out}$ | $P_{out}$ |
|---|---|---|---|---|---|---|
| *Combretum hereroense* | 0.96 | < 0.01* | 0.92 | < 0.01* | 0.97 | < 0.01* |
| *Dichrostachys cinerea* | 0.94 | < 0.01* | 0.96 | < 0.01* | 0.91 | < 0.01* |
| *Euclea undulata* | 0.86 | < 0.01* | 0.93 | < 0.01* | 0.97 | < 0.01* |
| *Flueggea virosa* | 0.88 | < 0.01* | 0.7 | < 0.01* | 0.86 | < 0.01* |
| *Grewia flava* | 0.86 | < 0.01* | 0.91 | < 0.01* | 0.81 | < 0.01* |
| *Grewia monticola* | 0.92 | < 0.01* | 0.94 | < 0.01* | 0.90 | < 0.01* |
| *Gymnosporia buxifolia* | 0.93 | < 0.01* | 0.86 | < 0.01* | 0.67 | < 0.01* |
| *Senegalia erubescens* | 0.95 | < 0.01* | 0.95 | < 0.01* | 0.96 | < 0.01* |
| *Senegalia mellifera* | 0.97 | < 0.01* | 0.71 | < 0.01* | 0.94 | < 0.01* |
| *Vachellia karroo* | 0.82 | < 0.01* | 0.89 | < 0.01* | 0.94 | < 0.01* |
| *Vachellia tortilis* | 0.95 | < 0.01* | 0.93 | < 0.01* | 0.95 | < 0.01* |
| *Ximenia americana* | 0.85 | < 0.01* | 0.96 | < 0.01* | 0.88 | < 0.01* |
| *Ziziphus mucronata* | 0.95 | < 0.01* | 0.95 | < 0.01* | 0.95 | < 0.01* |

* denotes significance to the level alpha = 0.05.

**Supporting information 3**: sample size of each dominant woody species sampled in Madikwe Game Reserve (2000-2001, “Old” condition; 2019, “New” condition) and in Barokologadi Communal Property (2019, “Out” condition).

| **Species** | $n_{old}$ | $n_{new}$ | $n_{out}$ |
|---|---|---|---|
| *Combretum hereroense* | 49 | 54 | 31 |
| *Dichrostachys cinerea* | 327 | 482 | 193 |
| *Euclea undulata* | 91 | 53 | 49 |
| *Flueggea virosa* | 63 | 54 | 17 |
| *Grewia flava* | 231 | 187 | 69 |
| *Grewia monticola* | 24 | 49 | 21 |
| *Gymnosporia buxifolia* | 57 | 35 | 25 |
| *Senegalia erubescens* | 261 | 131 | 38 |
| *Senegalia mellifera* | 23 | 15 | 67 |
| *Vachellia karroo* | 108 | 54 | 16 |
| *Vachellia tortilis* | 302 | 197 | 139 |
| *Ximenia americana* | 42 | 24 | 29 |
| *Ziziphus mucronata* | 111 | 56 | 18 |

**Supporting information 4**: Species-specific variations of tree height and canopy area according to spatio-temporal condition.

**Table S4.1**: Effect of spatio-temporal condition on the log-transformed tree height and canopy area in Madikwe Game Reserve (MGR) and Barokologadi Communal Property (BCP), South Africa: "Old", data collected in MGR in 2000 and 2001 at relatively low elephant density; "New", data collected in MGR in 2019 at relatively high elephant density; "Out", data collected in BCP in 2019 in the absence of elephants. The significance of the difference between two spatio-temporal conditions estimated with the linear mixed model were reported in the following columns: $P_{height\ new\text{-}old}$, $P_{height\ out\text{-}old}$, $P_{height\ new\text{-}out}$, $P_{area\ new\text{-}old}$, $P_{area\ out\text{-}old}$, $P_{area\ new\text{-}out}$.

| **Species** | $\mathbf{P_{height\ new\text{-}old}}$ | $\mathbf{P_{height\ out\text{-}old}}$ | $\mathbf{P_{height\ new\text{-}out}}$ | $\mathbf{P_{area\ new\text{-}old}}$ | $\mathbf{P_{area\ out\text{-}old}}$ | $\mathbf{P_{area\ new\text{-}out}}$ |
|---|---|---|---|---|---|---|
| *Combretum hereroense* | 0.34 | 0.01* | 0.02* | 0.11 | < 0.01* | 0.02* |
| *Dichrostachys cinerea* | < 0.01* | < 0.01* | < 0.01* | 0.12 | < 0.01* | < 0.01* |
| *Euclea undulata* | 0.36 | 0.01* | 0.04* | 0.10 | 0.01* | 0.22 |
| *Flueggea virosa* | < 0.01* | < 0.01* | 0.32 | < 0.01* | < 0.01* | 0.36 |
| *Grewia flava* | < 0.01* | 0.46 | 0.01* | 0.74 | 0.17 | 0.12 |
| *Grewia monticola* | 0.03* | 0.09 | < 0.01* | 0.79 | 0.04* | 0.02* |
| *Gymnosporia buxifolia* | 0.23 | 0.99 | 0.54 | 0.70 | 0.79 | 0.95 |
| *Senegalia erubescens* | 0.04* | 0.23 | 0.64 | 0.07 | 0.29 | 0.67 |
| *Senegalia mellifera* | 0.02* | 0.17 | 0.24 | 0.27 | 0.54 | 0.55 |
| *Vachellia karroo* | 0.05 | 0.01* | < 0.01* | 0.81 | < 0.01* | < 0.01* |
| *Vachellia tortilis* | < 0.01* | 0.02* | < 0.01* | < 0.01* | < 0.01* | < 0.01* |
| *Ximenia americana* | 0.03* | 0.66 | 0.55 | 0.64 | 0.87 | 0.96 |
| *Ziziphus mucronata* | < 0.01* | 0.25 | < 0.01* | < 0.01* | 0.11 | < 0.01* |

* denotes significance to the level alpha = 0.05.

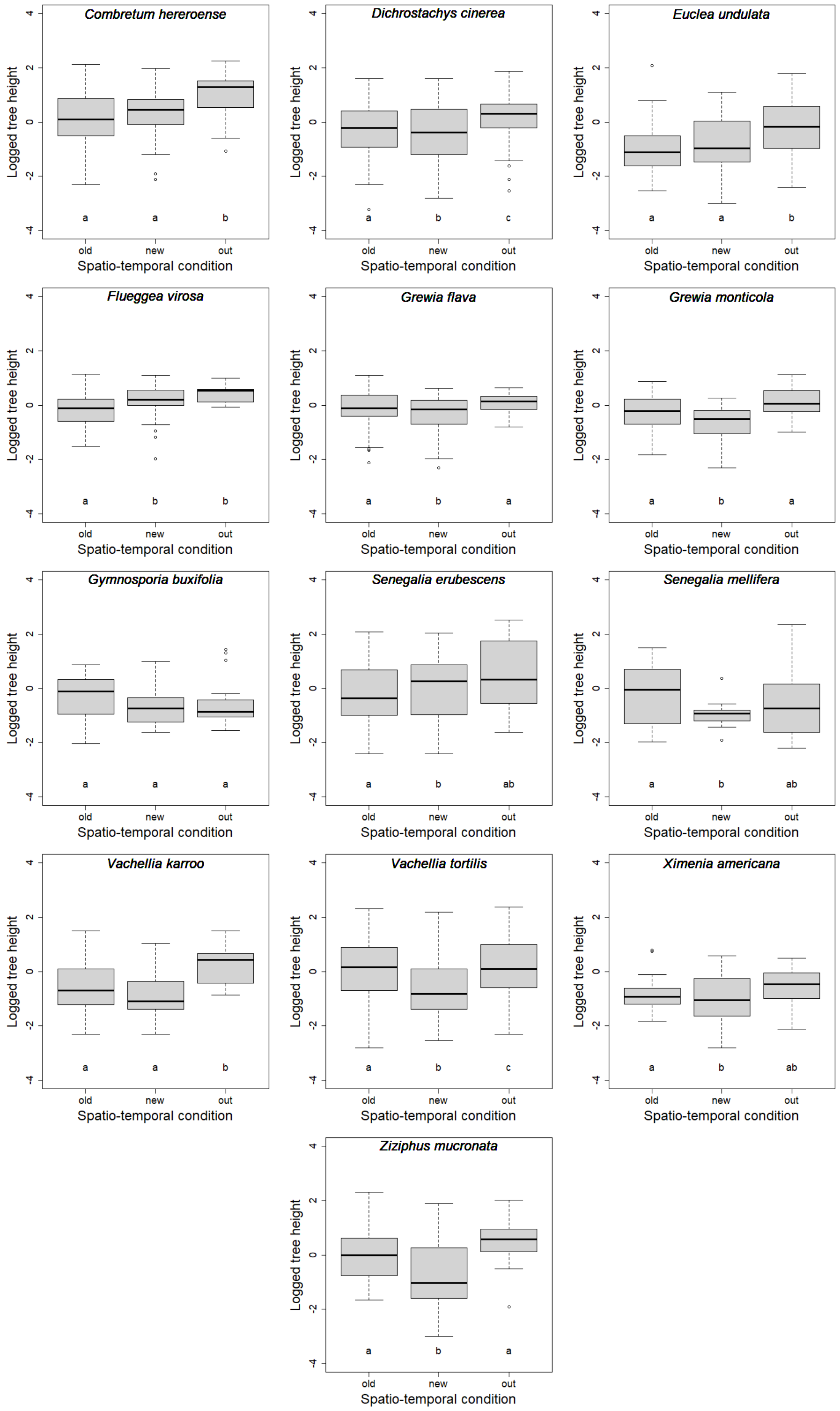


**Figure S4.1**: Species-specific variation of the logged tree height between the three spatio-temporal conditions: "Old", data collected in MGR in 2000 and 2001 at relatively low elephant density; "New", data collected in MGR in 2019 at relatively high elephant density; "Out", data collected in BCP in 2019 in the absence of elephants.*

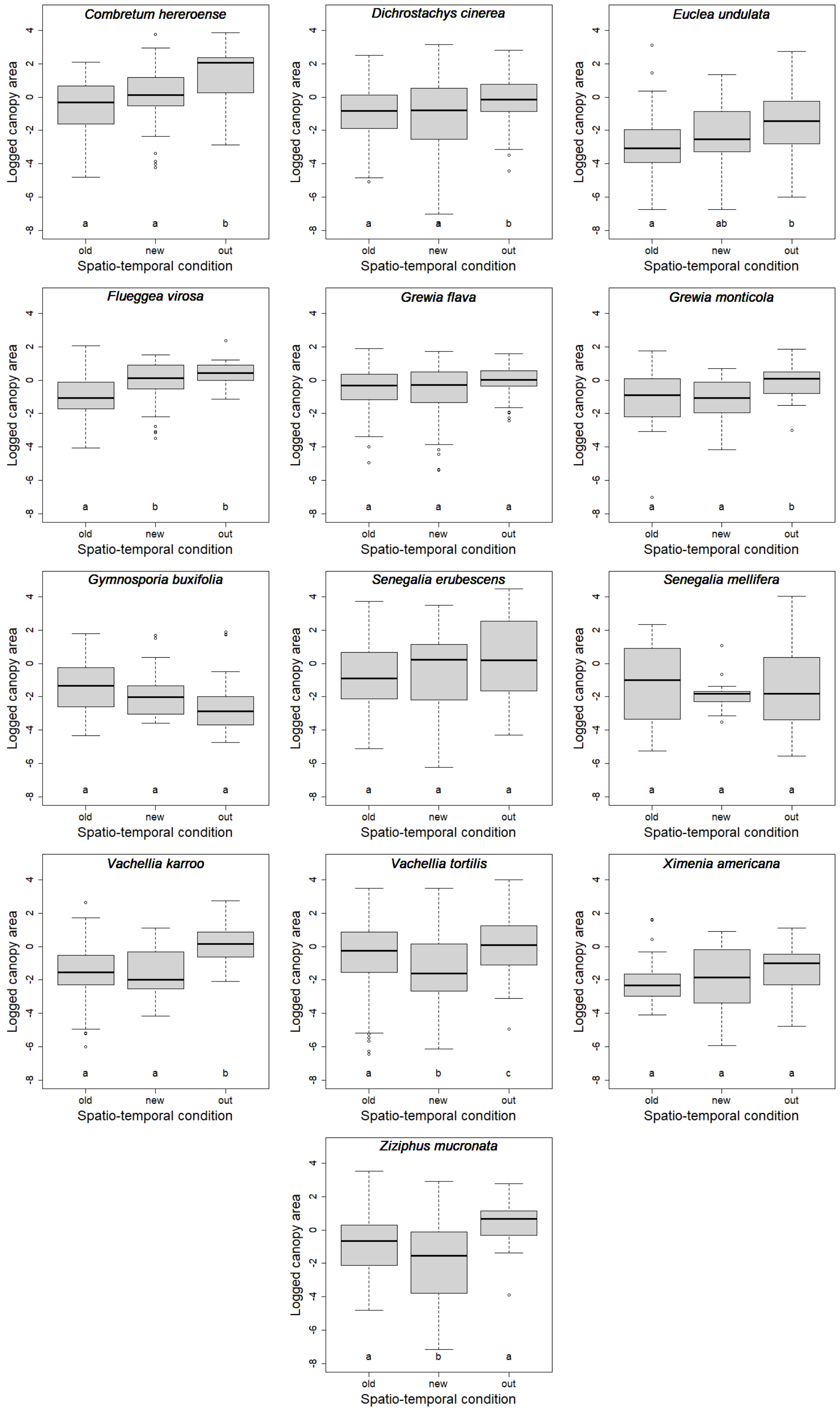


**Figure S4.2**: Species-specific variation of the logged canopy area between the three spatio-temporal conditions: “Old”, data collected in MGR in 2000 and 2001 at relatively low elephant density; “New”, data collected in MGR in 2019 at relatively high elephant density; “Out”, data collected in BCP in 2019 in the absence of elephants.*

*In the figures, black vertical lines correspond to the median, grey boxes correspond to the interquartile range (IQR) of the distribution, whiskers extend to the minimum and maximum values within 1.5 times the IQR and open dots correspond to the values beyond these limits.

**Supporting information 5**: Species-specific structural relationships.

In the following figures, the datasets were collected in Madikwe Game Reserve (2000-2001 and 2019, n = 35 sites) and in Barokologadi Communal Property (2019, n = 27 sites). The three spatio-temporal conditions are represented in: orange for the “Old” condition, red for the “New” condition, blue for the “Out” condition. Dots correspond to the raw observations, lines correspond to the predictions of the model, shaded areas represent the 95 % confidence intervals (calculated on the fixed effects). Note that the log scale (natural log, inside labels) and the real scale (outside labels) are provided on the graphs.

* Due to convergence problems, random effects (i.e., sampling site) were not included in six models for the tree height - stem diameter relationship (*Flueggea virosa* – New and Out, *Grewia monticola* – out, *Gymnosporia buxifolia* – Out, *Senegalia mellifera* – Old, *Ziziphus mucronata* – Out), six models for the canopy area - stem diameter relationship (*Flueggea virosa* – Out, *Grewia flava* – Out, *Grewia monticola* – Out, *Gymnosporia buxifolia* – Out, *Senegalia mellifera* – Old and Out) and nine models for the maximum vertical diameter - stem diameter relationship (*Euclea undulata* – New, *Flueggea virosa* – Out, *Grewia monticola* – Out, *Senegalia erubescens* – Old, *Senegalia mellifera* – Old, New and Out, *Vachellia karroo* – Out, *Ziziphus mucronata* – Out).

### Structural relationship between logged tree height and logged stem diameter

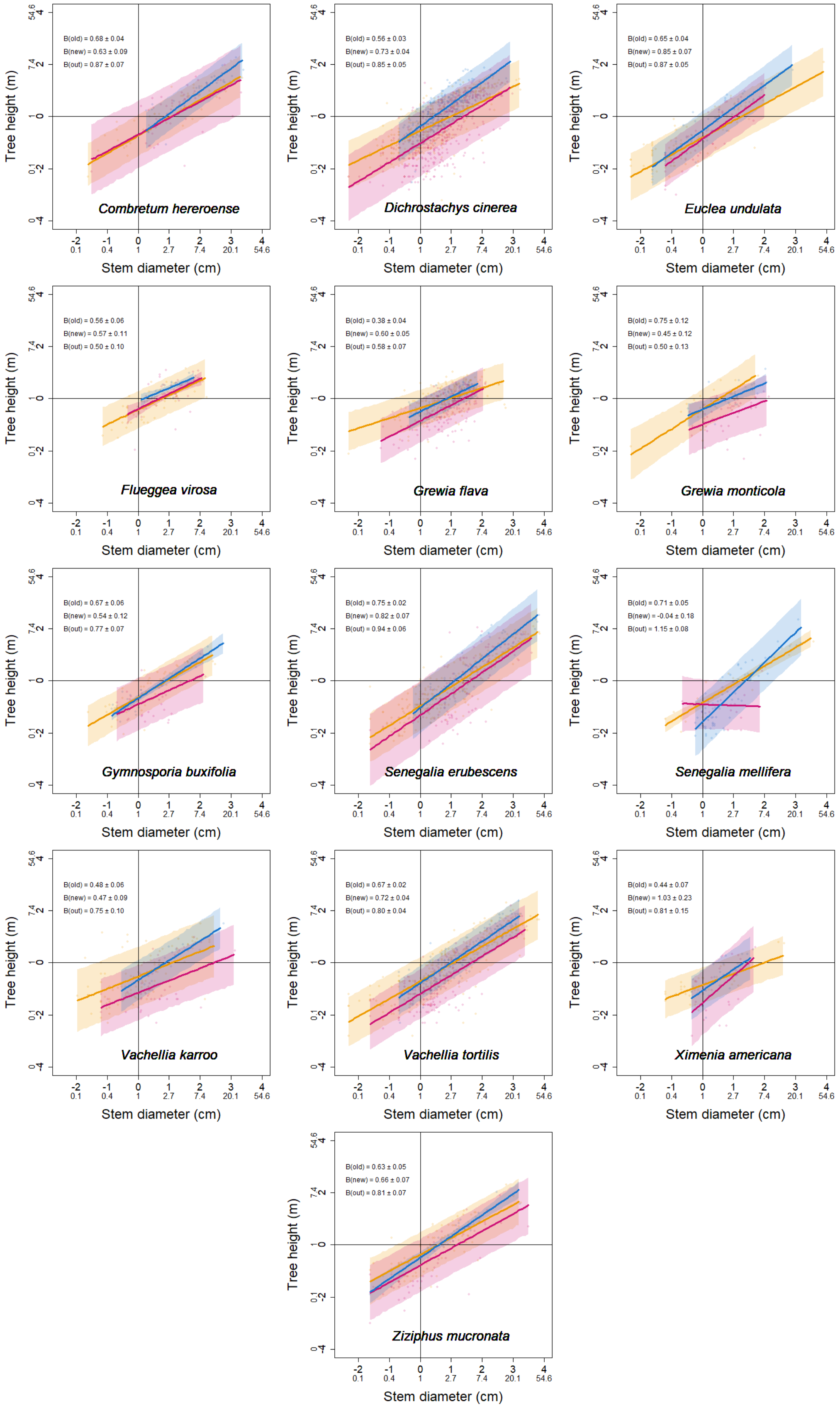

### Structural relationship between logged canopy vertical area and logged stem diameter

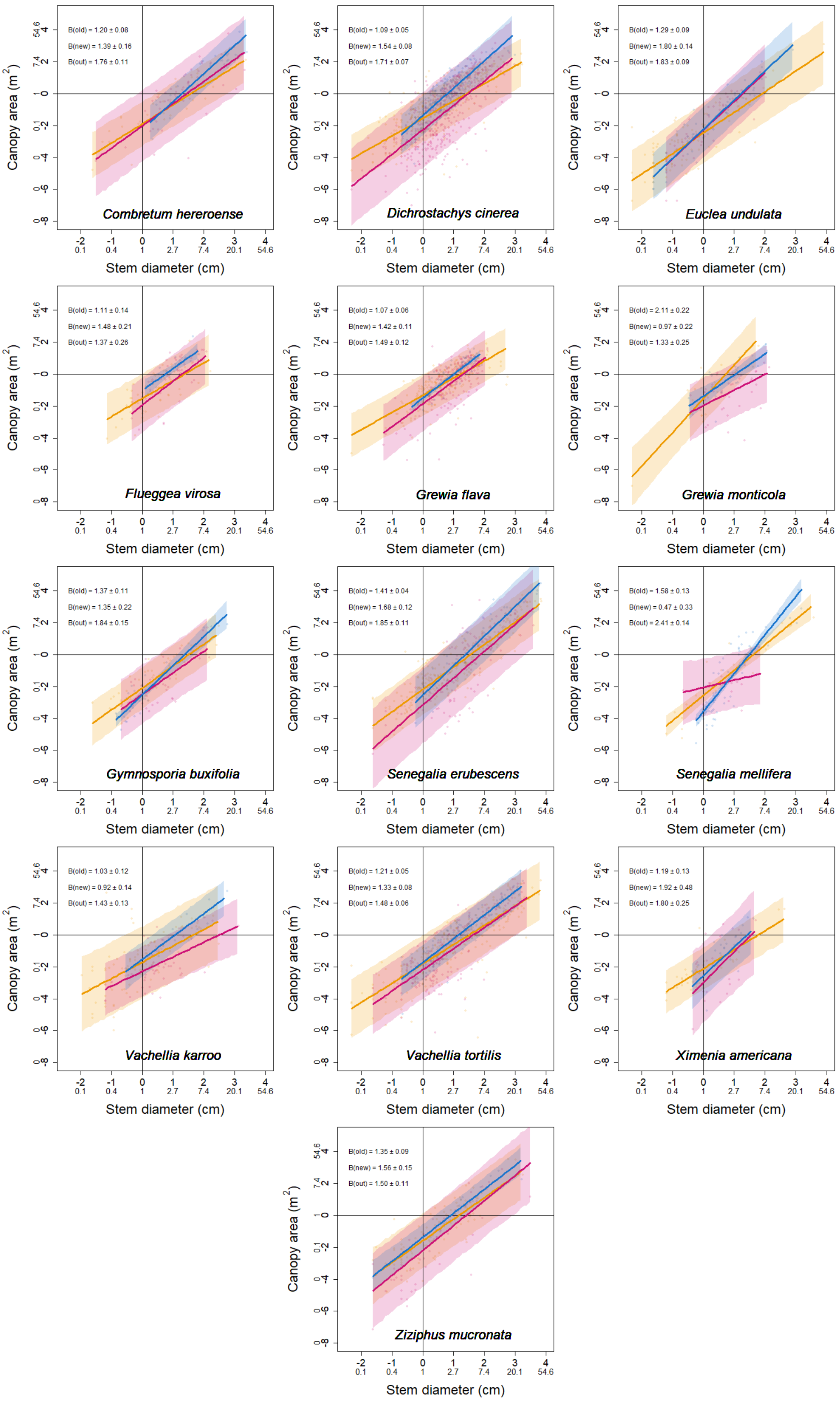

#### Structural relationship between logged max vertical diameter of the canopy and logged stem diameter

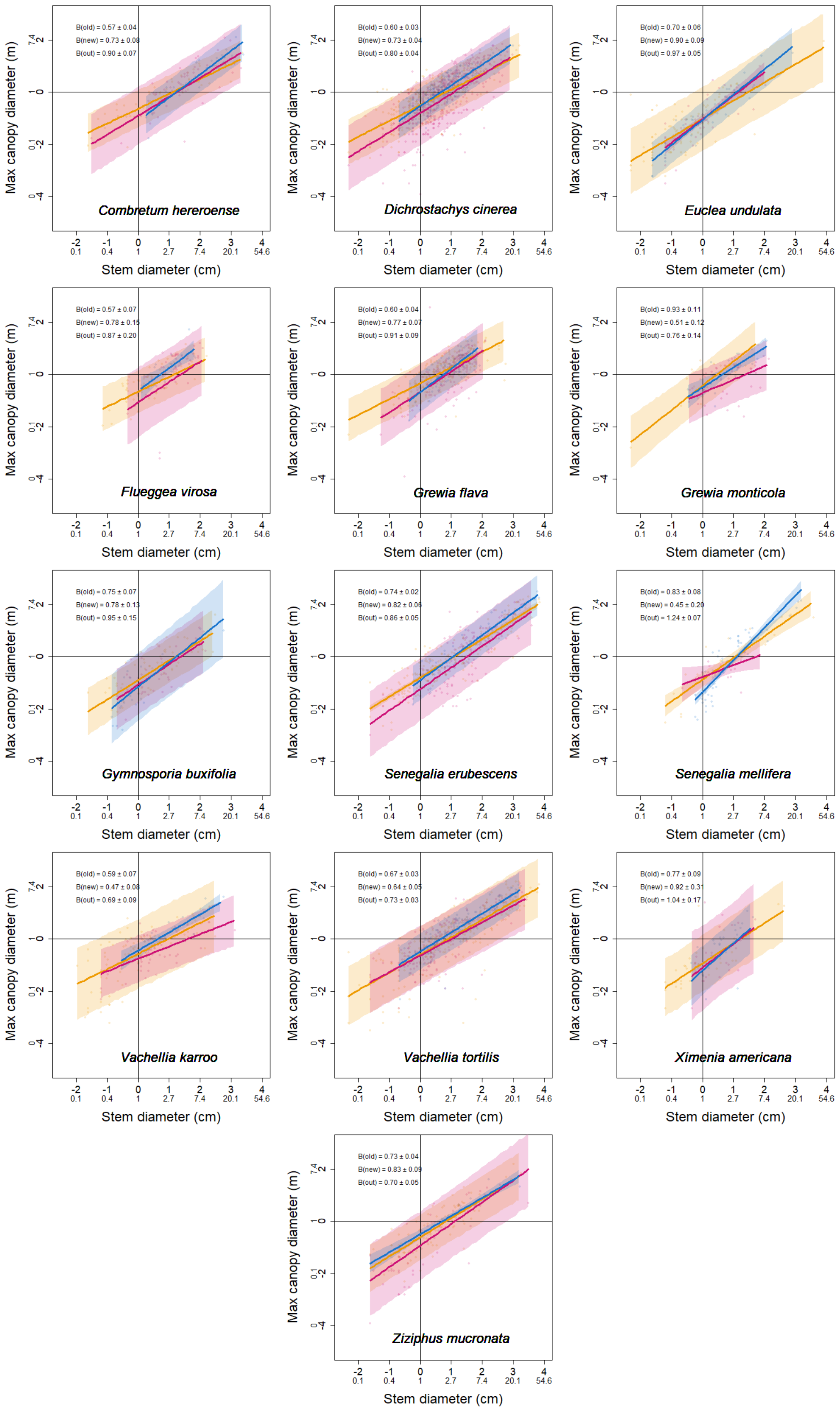

**Supporting information 6**: Residuals of the predictions of the structural relationships.

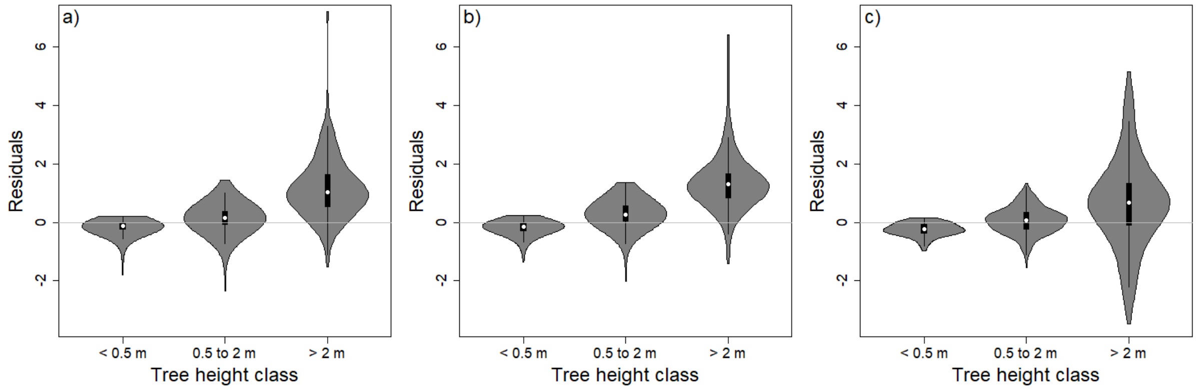


**Figure S6.1**: Distribution of the residuals of the fit of the structural relationship between tree height and stem diameter per tree height class (< 0.5 m, from 0.5 to 2 m, > 2 m) for the three spatio-temporal conditions: a) “Old”, data collected in MGR in 2000 and 2001 at relatively low elephant density; b) “New”, data collected in MGR in 2019 at relatively high elephant density; c) “Out”, data collected in BCP in 2019 in the absence of elephants.*

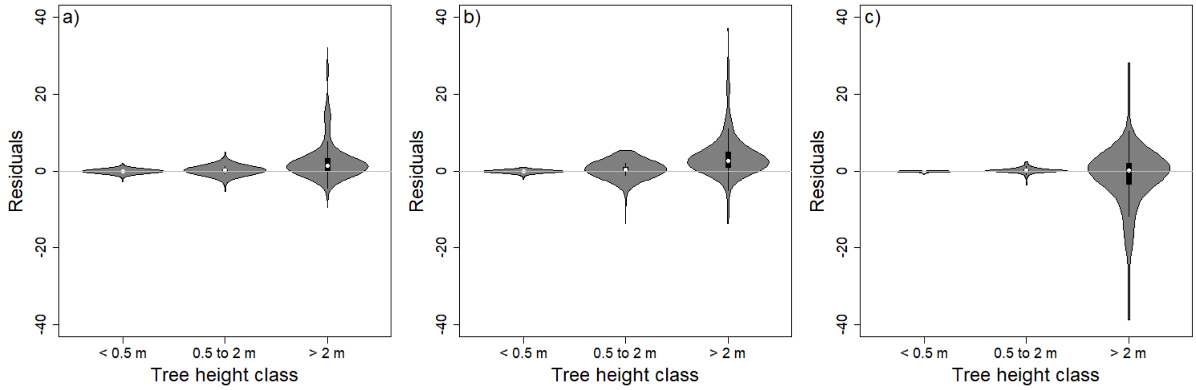


**Figure S6.2**: Distribution of the residuals of the fit of the structural relationship between canopy area and stem diameter per tree height class (< 0.5 m, from 0.5 to 2 m, > 2 m) for the three spatio-temporal conditions: a) “Old”, data collected in MGR in 2000 and 2001 at relatively low elephant density; b) “New”, data collected in MGR in 2019 at relatively high elephant density; c) “Out”, data collected in BCP in 2019 in the absence of elephants.*

*In the figures, white dots correspond to the median, black boxes correspond to the interquartile range (IQR) of the distribution, whiskers extend to the minimum and maximum values within 1.5 times the IQR and shaded areas correspond to the kernel density estimation of the distribution.

**Supporting information 7**: Elephant utilisation index according to tree height class.

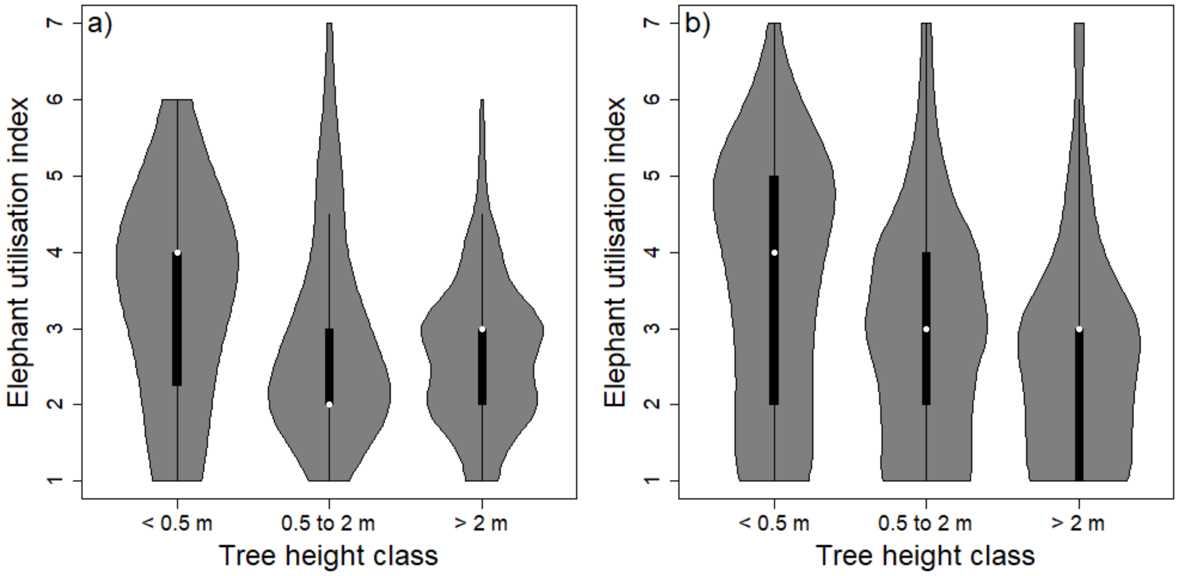


**Figure S7.1**: Distribution of elephant utilisation index per tree height class (< 0.5 m, from 0.5 to 2 m, > 2 m) in Madikwe Game Reserve: a) "Old", data collected in 2000 and 2001 at relatively low elephant density; b) "New", data collected in 2019 at relatively high elephant density. White dots correspond to the median, black boxes correspond to the interquartile range (IQR) of the distribution, whiskers extend to the minimum and maximum values within 1.5 times the IQR and shaded areas correspond to the kernel density estimation of the distribution.

**Supporting information 8**: Effect of spatio-temporal condition and tree height class on stem diameter and canopy area.

**Table S8.1**: Outputs of the linear mixed model describing the effect of the interaction between spatio-temporal condition and tree height class on stem diameter (with species and sampling site as random intercepts) in Madikwe Game Reserve (MGR) and Barokologadi Communal Property (BCP), South Africa. The three spatio-temporal conditions are: i) "Old", data collected in MGR in 2000 and 2001 at low elephant density; ii) "New", data collected in MGR in 2019 at high elephant density; iii) "Out", data collected in BCP in 2019 in the absence of elephants. The three tree height classes are: i) < 0.5 m, ii) from 0.5 to 2 m, iii) > 2 m.

| **Condition 1** | **Tree height class 1** | **Condition 2** | **Tree height class 2** | **beta ± standard error** | **P** |
|---|---|---|---|---|---|
| old | < 0.5 m | old | 0.5 to 2 m | 1.03 ± 0.04 | < 0.01* |
| old | < 0.5 m | old | > 2 m | 2.47 ± 0.05 | < 0.01* |
| old | < 0.5 m | new | < 0.5 m | 0.53 ± 0.04 | < 0.01* |
| old | < 0.5 m | new | 0.5 to 2 m | -0.19 ± 0.05 | < 0.01* |
| < 0.01* | < 0.5 m | new | > 2 m | -0.71 ± 0.08 | < 0.01* |
| < 0.01* | < 0.5 m | out | < 0.5 m | 0.30 ± 0.08 | < 0.01* |
| < 0.01* | < 0.5 m | out | 0.5 to 2 m | -0.18 ± 0.08 | 0.02* |
| old | < 0.5 m | out | > 2 m | -0.33 ± 0.09 | < 0.01* |
| old | 0.5 to 2 m | new | < 0.5 m | 0.19 ± 0.05 | < 0.01* |
| old | 0.5 to 2 m | new | 0.5 to 2 m | 0.34 ± 0.03 | < 0.01* |
| old | 0.5 to 2 m | new | > 2 m | -0.52 ± 0.07 | < 0.01* |
| old | 0.5 to 2 m | out | < 0.5 m | 0.18 ± 0.08 | 0.02* |
| old | 0.5 to 2 m | out | 0.5 to 2 m | 0.11 ± 0.06 | 0.08 |

| | | | | | |
|---|---|---|---|---|---|
| old | 0.5 to 2 m | out | > 2 m | -0.15 ± 0.08 | 0.06 |
| old | > 2 m | old | 0.5 to 2 m | -1.44 ± 0.04 | < 0.01* |
| old | > 2 m | new | < 0.5 m | 0.71 ± 0.08 | < 0.01* |
| old | > 2 m | new | 0.5 to 2 m | 0.52 ± 0.07 | < 0.01* |
| old | > 2 m | new | > 2 m | -0.18 ± 0.06 | < 0.01* |
| old | > 2 m | out | < 0.5 m | 0.33 ± 0.09 | < 0.01* |
| old | > 2 m | out | 0.5 to 2 m | 0.15 ± 0.08 | 0.06 |
| old | > 2 m | out | > 2 m | -0.04 ± 0.08 | 0.66 |
| new | < 0.5 m | new | 0.5 to 2 m | 0.84 ± 0.04 | < 0.01* |
| new | < 0.5 m | new | > 2 m | 1.76 ± 0.06 | < 0.01* |
| new | > 2 m | new | 0.5 to 2 m | -0.92 ± 0.06 | < 0.01* |
| out | < 0.5 m | new | < 0.5 m | 0.23 ± 0.08 | < 0.01* |
| out | < 0.5 m | new | 0.5 to 2 m | -0.01 ± 0.08 | 0.91 |
| out | < 0.5 m | new | > 2 m | -0.38 ± 0.10 | < 0.01* |
| out | < 0.5 m | out | 0.5 to 2 m | 0.85 ± 0.07 | < 0.01* |
| out | < 0.5 m | out | > 2 m | 2.14 ± 0.08 | < 0.01* |
| out | 0.5 to 2 m | new | < 0.5 m | 0.01 ± 0.08 | 0.91 |
| out | 0.5 to 2 m | new | 0.5 to 2 m | 0.22 ± 0.06 | < 0.01* |
| out | 0.5 to 2 m | new | > 2 m | -0.37 ± 0.08 | < 0.01* |
| out | > 2 m | new | < 0.5 m | 0.38 ± 0.10 | < 0.01* |
| out | > 2 m | new | 0.5 to 2 m | 0.37 ± 0.08 | < 0.01* |
| out | > 2 m | new | > 2 m | -0.15 ± 0.09 | 0.10 |
| out | > 2 m | out | 0.5 to 2 m | -1.29 ± 0.06 | < 0.01* |

* denotes significance to the level alpha = 0.05.

**Table S8.2**: Outputs of the linear mixed model describing the effect of the interaction between spatio-temporal condition and tree height class on canopy area (with species and sampling site as random intercepts) in Madikwe Game Reserve (MGR) and Barokologadi Communal Property (BCP), South Africa. The three spatio-temporal conditions are: i) "Old", data collected in MGR in 2000 and 2001 at low elephant density; ii) "New", data collected in MGR in 2019 at high elephant density; iii) "Out", data collected in BCP in 2019 in the absence of elephants. The three tree height classes are: i) < 0.5 m, ii) from 0.5 to 2 m, iii) > 2 m.

| **Condition 1** | **Tree height class 1** | **Condition 2** | **Tree height class 2** | **beta ± standard error** | **P** |
|---|---|---|---|---|---|
| old | < 0.5 m | old | 0.5 to 2 m | 2.01 ± 0.06 | < 0.01* |
| old | < 0.5 m | old | > 2 m | 4.11 ± 0.07 | < 0.01* |
| old | < 0.5 m | new | < 0.5 m | -0.11 ± 0.06 | 0.08 |
| old | < 0.5 m | new | 0.5 to 2 m | 0.56 ± 0.08 | < 0.01* |
| old | < 0.5 m | new | > 2 m | 0.25 ± 0.11 | 0.02* |
| old | < 0.5 m | out | < 0.5 m | 0.08 ± 0.11 | 0.50 |
| old | < 0.5 m | out | 0.5 to 2 m | 0.30 ± 0.11 | 0.01* |
| old | < 0.5 m | out | > 2 m | 0.42 ± 0.14 | < 0.01* |
| old | 0.5 to 2 m | new | < 0.5 m | -0.56 ± 0.08 | < 0.01* |
| old | 0.5 to 2 m | new | 0.5 to 2 m | 0.46 ± 0.05 | < 0.01* |
| old | 0.5 to 2 m | new | > 2 m | -0.31 ± 0.10 | < 0.01* |
| old | 0.5 to 2 m | out | < 0.5 m | -0.30 ± 0.11 | 0.01* |
| old | 0.5 to 2 m | out | 0.5 to 2 m | 0.38 ± 0.08 | < 0.01* |
| old | 0.5 to 2 m | out | > 2 m | 0.12 ± 0.11 | 0.28 |
| old | > 2 m | old | 0.5 to 2 m | -2.10 ± 0.07 | < 0.01* |
| old | > 2 m | new | < 0.5 m | -0.25 ± 0.11 | 0.02* |

| | | | | | |
|---|---|---|---|---|---|
| old | > 2 m | new | 0.5 to 2 m | 0.31 ± 0.10 | < 0.01* |
| old | > 2 m | new | > 2 m | 0.14 ± 0.09 | 0.12 |
| old | > 2 m | out | < 0.5 m | -0.42 ± 0.14 | < 0.01* |
| old | > 2 m | out | 0.5 to 2 m | -0.12 ± 0.11 | 0.28 |
| old | > 2 m | out | > 2 m | 0.50 ± 0.11 | < 0.01* |
| new | < 0.5 m | new | 0.5 to 2 m | 2.57 ± 0.06 | < 0.01* |
| new | < 0.5 m | new | > 2 m | 4.36 ± 0.09 | < 0.01* |
| new | > 2 m | new | 0.5 to 2 m | -1.79 ± 0.08 | < 0.01* |
| out | < 0.5 m | new | < 0.5 m | -0.18 ± 0.11 | 0.10 |
| out | < 0.5 m | new | 0.5 to 2 m | 0.26 ± 0.11 | 0.02* |
| out | < 0.5 m | new | > 2 m | -0.17 ± 0.14 | 0.23 |
| out | < 0.5 m | out | 0.5 to 2 m | 2.31 ± 0.10 | < 0.01* |
| out | < 0.5 m | out | > 2 m | 4.53 ± 0.12 | < 0.01* |
| out | 0.5 to 2 m | new | < 0.5 m | -0.26 ± 0.11 | 0.02* |
| out | 0.5 to 2 m | new | 0.5 to 2 m | 0.08 ± 0.08 | 0.36 |
| out | 0.5 to 2 m | new | > 2 m | -0.43 ± 0.12 | < 0.01* |
| out | > 2 m | new | < 0.5 m | 0.17 ± 0.14 | 0.23 |
| out | > 2 m | new | 0.5 to 2 m | 0.43 ± 0.12 | < 0.01* |
| out | > 2 m | new | > 2 m | -0.36 ± 0.12 | < 0.01* |
| out | > 2 m | out | 0.5 to 2 m | -2.22 ± 0.09 | < 0.01* |

* denotes significance to the level alpha = 0.05.
